# Age of onset in genetic prion disease and the design of preventive clinical trials

**DOI:** 10.1101/401406

**Authors:** Eric Vallabh Minikel, Sonia M Vallabh, Margaret C Orseth, Jean-Philippe Brandel, Stéphane Haïk, Jean-Louis Laplanche, Inga Zerr, Piero Parchi, Sabina Capellari, Jiri Safar, Janna Kenny, Jamie C Fong, Leonel T Takada, Claudia Ponto, Peter Hermann, Tobias Knipper, Christiane Stehmann, Tetsuyuki Kitamoto, Ryusuke Ae, Tsuyoshi Hamaguchi, Nobuo Sanjo, Tadashi Tsukamoto, Hidehiro Mizusawa, Steven J Collins, Roberto Chiesa, Ignazio Roiter, Jesús de Pedro-Cuesta, Miguel Calero, Michael D Geschwind, Masahito Yamada, Yosikazu Nakamura, Simon Mead

## Abstract

Regulatory agencies worldwide have adopted programs to facilitate drug development for diseases where the traditional approach of a randomized trial with a clinical endpoint is expected to be prohibitively lengthy or difficult. Here we provide quantitative evidence that this criterion is met for the prevention of genetic prion disease. We assemble age of onset or death data from *N*=1,094 individuals with high penetrance mutations in the prion protein gene (*PRNP*), generate survival and hazard curves, and estimate statistical power for clinical trials. We show that, due to dramatic and unexplained variability in age of onset, randomized preventive trials would require hundreds or thousands of at-risk individuals in order to be statistically powered for an endpoint of clinical onset, posing prohibitive cost and delay and likely exceeding the number of individuals available for such trials. Instead, the characterization of biomarkers suitable to serve as surrogate endpoints will be essential for the prevention of genetic prion disease. Biomarker-based trials may require post-marketing studies to confirm clinical benefit. Parameters such as longer trial duration, increased enrollment, and the use of historical controls in a post-marketing study could provide opportunities for subsequent determination of clinical benefit.

## Introduction

Placebo-controlled, double-blind, randomized trials with a clinical endpoint — a measure of how patients feel or function — constitute the gold standard for demonstration of therapeutic efficacy and, where feasible, are strongly preferred for approval of new drugs. Regulators worldwide have recognized, however, that in some diseases the duration of such trials may unduly delay patient access to potentially life-saving drugs. Many agencies have therefore created programs to support drug development in this situation. For instance, the United States Food and Drug Administration (FDA) Accelerated Approval program^1^ provides for conditional approval based on trials using surrogate endpoints, including biomarkers, with a requirement for post-marketing studies to confirm clinical benefit^2^. Honoring the specifics of each disease, FDA “consider[s] how to incorporate novel approaches into the review of surrogate endpoints… especially in instances where the low prevalence of a disease renders the existence or collection of other types of data unlikely or impractical”^3^. Here we present evidence that genetic prion disease meets this criterion.

Prion disease is a fatal and, at present, incurable neurodegenerative disease caused by the misfolding of the prion protein, PrP, encoded by the gene *PRNP*^4^. Most subtypes of prion disease are extremely rapid, leading from first symptom to death in several months^5^. Prion diseases are transmissible, but today few cases are known to be acquired by infection. ∼85% of prion disease cases are termed “sporadic,” meaning they arise spontaneously with no known environmental or genetic trigger, while ∼15% of cases possess protein-altering rare variants in *PRNP*, a subset of which are highly penetrant^6^. Various genetic subtypes of prion disease include fatal familial insomnia, genetic Creutzfeldt-Jakob disease, and Gertsmann-Sträussler-Scheinker disease.

To date, all completed clinical trials in prion disease have recruited only symptomatic patients, mostly with sporadic prion disease, and have used cognitive, functional, or survival endpoints^7^– ^14^. By the time of diagnosis many prion disease patients are in a state of advanced dementia, and even a therapy that halted the disease process entirely at this stage might only preserve the patient in a state with little or no quality of life^15^. Moreover, preclinical proofs of concept argue that a preventive, rather than therapeutic, approach is more likely to be effective. Multiple antiprion agents have been discovered that extend the survival time of prion-infected mice by 2-4X when administered long before symptoms, yet these have diminished effects at later timepoints, and no effect when administered after clinical onset^16–19^. These observations indicate a need to enable preventive trials in presymptomatic individuals at risk for genetic prion disease.

The ongoing preventive trial of crenezumab, an anti-amyloid β antibody, for *PSEN1* E280A early-onset Alzheimer’s disease, follows a design where presymptomatic individuals are randomized to drug or placebo and followed for five years to a cognitive endpoint^20^. While this represents one model for preventive trials in neurodegeneration, we hypothesized that this approach might be challenging for genetic prion disease due to its variable age of onset^21,22^, small presymptomatic patient population^23^, and more limited financial incentives for pharmaceutical companies. To test this hypothesis, we set out to aggregate age of onset data in genetic prion disease, generate survival and hazard curves, and simulate statistical power for randomized pre-approval trials with a clinical endpoint. We also set out to investigate the feasibility of one potential alternative: post-marketing studies using historical controls to confirm clinical benefit, following Accelerated Approval on a surrogate biomarker endpoint.

## Results

### Age of onset in genetic prion disease

We reasoned that any preventive trial with a clinical endpoint in genetic prion disease would derive most of its statistical power from individuals with high penetrance *PRNP* variants. Some *PRNP* variants can be identified as highly penetrant by their extreme enrichment in cases over population controls, but many variants are too rare in both groups for meaningful comparison^6^. We therefore reviewed the literature on 69 reportedly pathogenic *PRNP* variants and identified 27 variants for which Mendelian segregation has been reported in at least one family with at least three affected individuals and/or for which a *de novo* mutation in a case has been identified (Table S1), thus suggesting high penetrance.

We examined the frequency of these putative high penetrance variants in a recent case series^6^. The top three variants — E200K, P102L, and D178N — collectively explain 85% of high penetrance cases (Figure S1). Each of these arises from a CpG transition (a C to T DNA change where the adjacent base is G), a type of variant which occurs by spontaneous mutation 10-100X more often than other mutation types^24,25^, explaining the recurrence of these three mutations on multiple *PRNP* haplotypes in families worldwide^6,26,27^. Therefore, regardless of the population studied, these three variants are likely to account for a large fraction of genetic prion disease cases with high penetrance variants. For this reason, we focused our analysis primarily on individuals with these three variants. We aggregated age of onset and/or age of death data on *N*=1,001 individuals with the E200K, P102L, or D178N mutations from nine study centers worldwide (Table 1 and Table S2), encompassing both direct clinical reports and family histories (see Methods), and including censored individuals. Statistics on *N*=93 individuals with the next four mutations most common in cases — 5-OPRI (insertion of five extra octapeptide repeats), 6-OPRI, P105L, and A117V — are included in Table S3. We used these data to compile life tables and computed the annual hazard — risk of onset in each year of life — for each mutation (Supplementary Life Tables).

We found wide variability in age of onset (Table 1), consistent with previous reports^21,22,28^. An implication of this variability is that high lifetime risk arises not from certain onset at a specific age, but from modest risk in any given year of life, accumulated over many decades of exposure. This poses a challenge for following presymptomatic individuals to onset in a preventive clinical trial, as it is difficult to ascertain a group of individuals for whom onset is imminent. For example, even at age 57, an E200K individual has only a 5% probability of disease onset occurring in any given year. This means that 20 person-years of follow-up for E200K individuals around this age would be expected to result in only one observed disease onset. Annual hazards do rise with age, but as they reach high levels, the number of surviving individuals also dwindles (Figure 1). For the three most common mutations, the annual hazard remains below 10% until after the majority of people have already died (Figure 1, Figure S3, and Supplementary Life Tables). Similarly, the median number of years until onset, conditioned on an individual’s current age, remains ≥5 years until after the median age of onset has passed (Supplementary Life Tables). The next four most common mutations have tighter age of onset distributions, and so reach higher annual hazards sooner (Figure S3), but these mutations are also much rarer, accounting for only 10% of cases with a high penetrance variant^6^.

**Table 1.** Variability in age of onset in genetic prion disease. Censored data include individuals who were either alive and well at last follow-up or had died of an unrelated cause, and whose genetic status was known either through predictive testing or due to obligate carrier status. For D178N and E200K, because the majority of individuals have disease duration ≤1 year (Figure S2), age of death is used where age of onset is unavailable. For P102L, which more often has a longer disease duration, only age of onset is used. IQR, interquartile range. Range indicates highest and lowest observed age of onset, except where * indicates that the longest survival is a censored data point.

| <u>mutation</u> | <u>without censored data</u> |  | <u>survival curve including censored data</u> |  |  |
| --- | --- | --- | --- | --- | --- |
|  | <u>mean ± sd</u> | <u>N</u> | <u>median (IQR)</u> | <u>range</u> | <u>N</u> |
| P102L | 53.7 ± 10.6 | 193 | 56 (47 - 60) | 22 - 75 | 206 |
| D178N | 51.3 ± 11.8 | 256 | 53 (46 - 60) | 12 - 89* | 289 |
| E200K | 61.3 ± 10.0 | 456 | 62 (55 - 68) | 31 - 92 | 506 |

**Figure 1.**
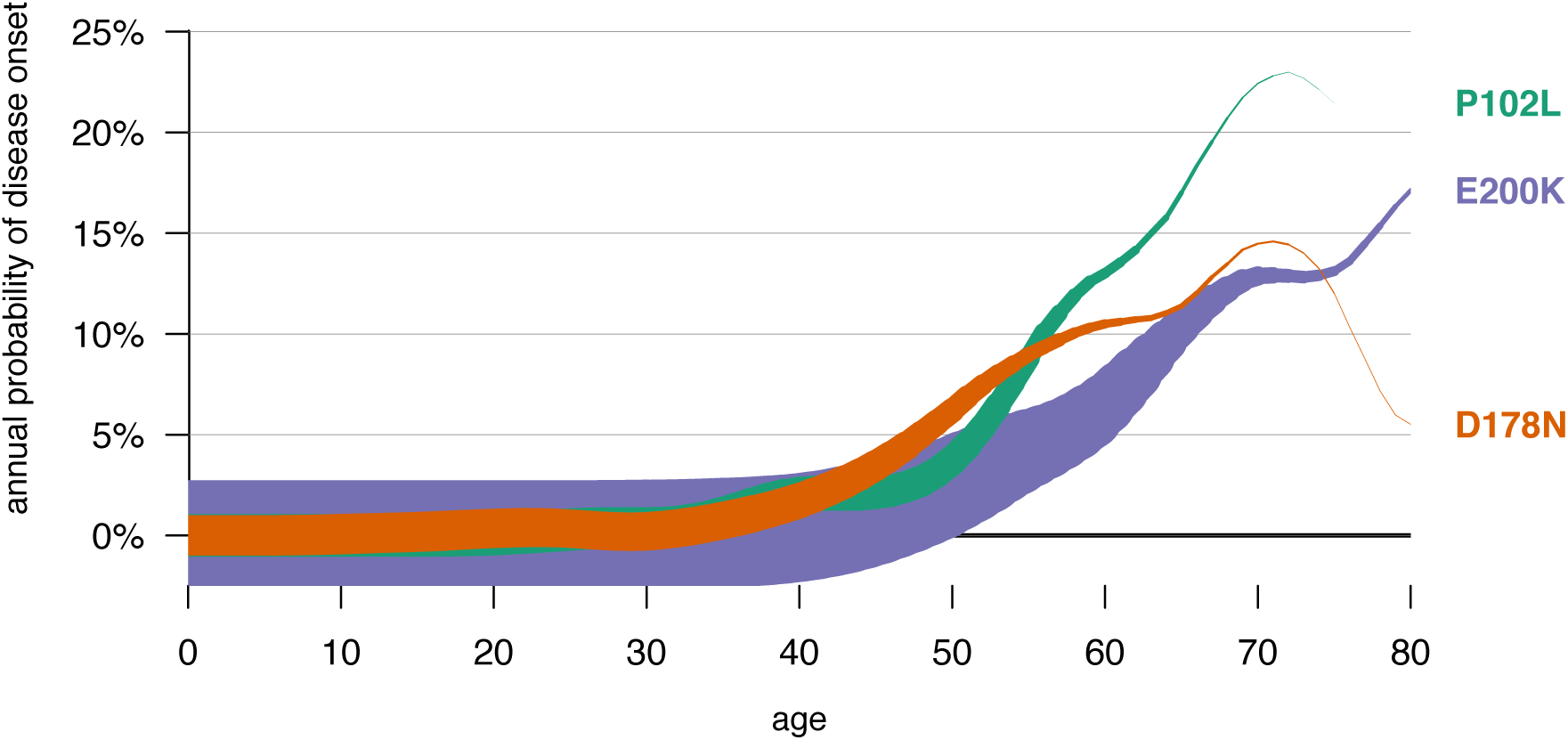
Hazards and survival for the most common PRNP mutations. The hazard, or probability of disease onset, in each year of life (y axis) is plotted against age (x axis) with curve thickness representing the number of individuals still living at each age, which is the product of age-dependent survival and mutation prevalence. Supplementary mutations, and conventional survival curves and hazard plots, are included in Figure S3.

### Power for randomized pre-approval trials with a clinical endpoint

We set out to calculate how many individuals would need to enroll in order to power prevention trials with an endpoint of disease onset, using the calculated age-dependent hazards for each mutation. While younger individuals or those with a mutation of modest penetrance might seek to enroll in trials or take a preventive drug, they would not contribute much statistical power to an endpoint of clinical onset. We therefore chose to base our power calculations on individuals with the three most common high penetrance mutations between age 40 and 80.

We estimated how many individuals in this age range have high penetrance *PRNP* mutations. It is estimated based on disease prevalence that 1-2 people per 100,000 in the general population harbor high penetrance *PRNP* mutations (ref. ^6^ and Supplementary Discussion), but at present, many remain unaware of their risk due to underdiagnosis^29^ of affected family members, and few choose predictive testing^23^ as the results are currently considered medically unactionable. The number of positive predictive genetic test results that have been provided in the U.S. is *N*=221 (Supplementary Discussion), and based on the estimated proportion of high penetrance variants^6^ (75%), and the estimated proportion of positive test result recipients^23^ over age 40 (36%) we estimate there are currently ∼60 people in the U.S. who are age 40 or older and hold a positive predictive test for a highly penetrant variant.

We used published formulae^30^ (see Methods) to calculate statistical power for a log-rank survival test in randomized clinical trials (Table 2). Across the three mutations and weighted by their prevalence among cases (Figure S1) and number of surviving individuals at each age (Figure 1), the average annual probability of onset for individuals aged 40 to 80 is 4.6%. We used the 4.6% figure as a baseline hazard, and made the following assumptions: pre-symptomatic individuals are randomized half to drug and half to placebo and followed for 5 years with an endpoint of clinical onset; events in the first year are ignored as a “run-in” period to ensure sufficient drug exposure among individuals analyzed; the withdrawal rate is 15.2% annually (the median value from eight prevention trials reviewed, Table S4); and the trial is designed for 80% power at the *P*=0.05 threshold. We then performed power calculations for such a trial as a function of the hazard ratio — the ratio of annual risk of onset in drug-treated individuals to that in placebo-treated individuals. For context, we also determined the effect size, in median years of healthy life added, to which each hazard ratio corresponds (Table 2). The calculations are sensitive to which mutations are included, the “run-in” period, the number of years of follow-up, and the assumed withdrawal rate, but we explored a range of different assumptions and none support a different overall interpretation of the data (Table S5 and Discussion). In particular, the assumption of a 15.2% annual withdrawal rate means that only 44% of original participants remain after 5 years, but even reducing the withdrawal rate to zero only lowers the numbers of participants required by one-third (Table S5). Because FDA has cautioned against rare disease trial designs that assume a large effect size^31^, we focus below on the moderate hazard ratio of 0.5, which would correspond to seven years of life added for treated individuals.

**Table 2.** Preventive trial requirements under survival test power calculation. For example, a hazard ratio of 0.5 means that placebo-treated individuals have a 4.6% annual probability of onset, while drug-treated individuals have only a 2.3% annual probability of onset. If a population of individuals were treated from an early age with such a drug, the median age of onset would be postponed by 7 years. To have 80% power at P=.05 to detect the effect of such a drug, 65 individuals would need to become symptomatic during the trial — given the 0.5 hazard ratio, about two-thirds of these would occur in the placebo group and one-third would occur in the drug group. Observing this number of disease onsets would require randomizing 813 people for 5 years (data from the first year would be ignored, and the remaining four years of data would be analyzed). *For a hazard ratio of 0.1, most individuals never become sick, thus, the increase in median age of onset (the age where 50% of people have had onset) is undefined.

| **hazard ratio** | **years of life added** | **onsets required** | **participants required** |
| --- | --- | --- | --- |
| 0.1 | undefined* | 6 | 101 |
| 0.2 | 21 | 12 | 189 |
| 0.3 | 14 | 22 | 311 |
| 0.4 | 10 | 37 | 498 |
| **0.5** | **7** | **65** | **813** |
| 0.6 | 5 | 120 | 1,406 |
| 0.7 | 4 | 247 | 2,724 |
| 0.8 | 2 | 631 | 6,602 |
| 0.9 | 1 | 2,828 | 28,204 |

The above power calculations simplistically assume a uniform baseline hazard across all participants, regardless of age and *PRNP* mutation. We also used a simulation to account for the full shape of the hazard curve and diversity of genetic mutations, but the simulated power results were similar to those in Table 2 (see Table S6). Stratification by *PRNP* mutation did not improve power in our simulations (Supplementary Discussion), perhaps because age of onset distributions (Table 1) are wide and overlapping, such that *PRNP* mutation explains only a minority of the overall variance in age of onset (adjusted R^2 = 0.15, linear regression, *P* < 1e-32).

Statistical power might be improved by stratifying clinical trial analysis by relevant additional variables, but there are currently no variables that help to explain age of onset (Supplementary Discussion, Table S7, and Figure S4-5). For instance, we found no sex effect, and no evidence that parent and child age of onset are correlated after controlling for *PRNP* mutation and for child’s year of birth, a variable that captures some effects of ascertainment bias^32^ (Table S7). A common genetic variant, *PRNP* M129V, is known to affect the clinical and pathological presentation of many forms of prion disease^33^ as well as the risk of sporadic and acquired prion disease^34^. This variant has previously been reported to affect age of onset in some forms of genetic prion disease but not others^21,32,35,36^. We found no evidence that codon 129 affects age of onset for P102L or E200K individuals (Table S7 and Figure S4). For D178N, our data are suggestive that a 129VV genotype may predispose to earlier onset than MM or MV genotypes (Figure S4 and Supplementary Discussion), but in the overall dataset, codon 129 failed to explain additional variance in age of onset (Supplementary Discussion).

Based on this analysis, at present it is not possible to adequately power a randomized pre-approval prevention trial with an endpoint of clinical onset in genetic prion disease. For example, for a drug that reduces annual risk by half (hazard ratio of 0.5), powering such a trial would require 813 participants age 40 or older, and even for a drug that reduces annual risk by ten-fold (hazard ratio of 0.1), 101 participants would be required (Table 2), versus the ∼60 currently estimated to exist in the U.S. Key assumptions underpinning this analysis may change with time: new stratifying variables could help to predict age of onset, or a first drug for prion disease could improve diagnosis and recruitment (see Discussion). However, the insight that randomized pre-approval prevention trials with a clinical endpoint may not be feasible today has implications for drug development efforts likely to reach the clinic while current assumptions hold. For this reason, we next turned our attention to the possibility that a preventive drug might be developed through the Accelerated Approval pathway using a surrogate biomarker endpoint.

### Power for post-marketing studies

We asked whether, if Accelerated Approval were achieved, the required post-marketing studies to confirm clinical benefit could be adequately powered by following drug-treated individuals to clinical onset and comparing their survival to that of historical controls. Such a trial design could increase power but also introduce bias; we considered each issue in turn.

We identified several factors that are likely to decrease the number of participants required to power such a study compared to its randomized pre-approval equivalent: all, rather than half, of individuals are drug-treated; the number of historical controls can be large; a longer trial duration could be considered because the trial would overlap, rather than reduce, the drug’s effective market exclusivity period (Supplementary Discussion); and a post-marketing surveillance program might allow newly drug-treated individuals to enter the program on a rolling basis, replacing any who withdraw. The effects of these assumptions (Table 3) are collectively to reduce the number of individuals required to demonstrate efficacy of a drug with hazard ratio of 0.5 from 813 to 37. At the same time, the number of individuals available for a trial might increase, because: an approved drug should have broader geographic reach than a pre-approval trial; a treatment might improve awareness and diagnosis of the disease; and a treatment might stimulate more individuals to pursue predictive genetic testing. For instance, of people at 50/50 risk for a *PRNP* mutation, currently only 23% pursue predictive testing, compared to 60% (2.6X higher) for *BRCA1* or *BRCA2* mutations^37^, which are considered medically actionable^38^. Thus, a post-marketing study could be adequately powered with available numbers of individuals for a hazard ratio of 0.5 (Table 3) and, over a range of assumptions, would bring power requirements into closer alignment with the number of available individuals (Figure S6 and Supplementary Discussion).

**Table 3.** Comparison of power calculations for pre-approval and post-marketing studies. In each case the calculation is for a hazard ratio of 0.5, the N indicated is for 80% power at the P=0.05 threshold, and all assumptions other than those indicated in the table are the same as for Table 2. The number of individuals required for post-marketing studies is determined by simulation (Supplementary Discussion).

| scenario | N required | explanation |
| --- | --- | --- |
| Randomized pre-approval — 5-year follow-up | 813 | See Table 2 |
| Post-marketing with historical controls — 5-year follow-up | 229 | Increased power because all, rather than half, of individuals are treated, and $N=1,000$ historical controls are used for comparison. |
| Post-marketing with historical controls — 15-year follow-up | 125 | Increased power because longer trial duration may be financially tenable for post-marketing studies. Power is still limited, however, by withdrawal rate, which means that few participants remain at the end of 15 years. |
| Post-marketing with historical controls — 15-year follow-up, no withdrawal | 37 | Increased power because the withdrawal rate is set to zero, simulating a scenario where individuals who go on drug can continuously enter the surveillance program, and the cohort being monitored can maintain its size over time. |

While we conducted tests to ensure that our power simulation was not itself biased (Supplementary Discussion), a post-marketing study could still be biased in real life, if the historical controls used do not accurately estimate the true hazard rates facing the trial participants^39^. There are no environmental, demographic, or non-*PRNP* genetic factors known to affect prion disease risk or age of onset, although these might nonetheless exist^40,41^. Perhaps of greater concern is that most of our historical controls were collected retrospectively — individuals are only ascertained if they become sick — and may overestimate the hazard rates for individuals followed prospectively^32^. To assess this possible source of bias, we compared the survival of the limited number of individuals followed prospectively in our dataset (*N*=24 individuals, with a cumulative 145 person-years of follow-up), conditioned on their ages at first ascertainment, to those of individuals with no prospective follow-up. We did not observe a significant difference in hazard (*P*=0.59, Cox proportional hazards test) between these two groups, although this could be due to a lack of power (see Discussion).

## Discussion

Both our power calculation and simulation indicate that direct demonstration of clinical benefit in a randomized pre-approval prevention trial would require enrolling a number of *PRNP* mutation carriers that is not currently realistic. For instance, for a drug that reduces annual risk of onset by half (hazard ratio of 0.5), estimated to correspond to a 7-year delay in median age of onset, 80% power was reached only with 813 individuals randomized for 5 years (Table 2). Currently, only *N*=221 presymptomatic individuals in the U.S. have positive genetic test results for *PRNP* mutations, and we estimate that only ∼60 of these have high penetrance mutations and fall in an age range (≥40 years) where their hazard is sufficiently high that they would contribute appreciable power to a randomized trial with a clinical endpoint. Randomized prevention trials might just barely achieve 80% power under the most wildly optimistic assumptions of an extremely effective drug (hazard ratio of 0.1, reducing annual risk of onset by ten-fold), along with some increase in predictive testing rates and a very successful trial recruitment effort. FDA has cautioned, however, that rare disease trials should not be designed around the hope of a huge effect size^31^, and even if a drug were so profoundly effective, it is unlikely that a sponsor would have sufficient confidence in this *a priori* to invest in a trial that is underpowered for more moderate effect sizes.

At least three factors can explain why a randomized trial design following pre-symptomatic individuals to a clinical endpoint was deemed feasible for early-onset Alzheimer’s disease, yet appears unviable for genetic prion disease. First, onset is less predictable in genetic prion disease. The standard deviation of age of onset ranges from ±10.0 to ±11.8 years for the three *PRNP* mutations we examined (Table 1), whereas estimates of the standard deviation of age of onset for *PSEN1* E280A Alzheimer’s disease range from ±6.4 to ±8.6 years^42,43^. In addition, an individual’s age of onset in genetic Alzheimer’s disease is reported to be correlated with parental age of onset^43^, and this property has been used to attempt to enrich for high-hazard individuals in trials^44^, whereas we have found no evidence that parent and child age of onset are correlated in genetic prion disease (Table S7 and ref. ^32^). Second, genetic prion disease is rarer. The *PSEN1* preventive trial recruited from a single pedigree of ∼5,000 individuals^45^ from which 1,065 living individuals with the mutation have been enrolled in a registry^46^. There is no known genetic prion disease family this large. Third, genetic prion disease offers more limited financial incentives for a pharmaceutical sponsor. The cost of the *PSEN1* preventive trial has been estimated at $96 million^44,47^, and while this price may be tenable for sponsors in view of potential for an expanded Alzheimer’s indication, no similar potential exists for prion disease. Indeed, even in Alzheimer’s disease, larger or longer primary prevention trials are likely to prove challenging for the private sector and may require public sector investment^48^.

Preclinical proof-of-concept studies in mice have shown that some antiprion agents effective at delaying prion disease on a prophylactic basis become ineffective if given close to the time of clinical onset^16,19^, suggesting that trials in symptomatic patients could fail to show a benefit that would have been realizable in preventive treatment. Yet our results here indicate that it would be difficult or impossible to design a well-powered randomized preventive trial with a clinical endpoint in genetic prion disease. Together, these observations argue for the characterization of biomarkers suitable as endpoints in presymptomatic genetic prion disease, and for their evaluation by regulatory agencies as surrogate trial endpoints. Accompanying manuscripts describe one possible route to Accelerated Approval using a surrogate biomarker endpoint^49,50^.

If Accelerated Approval could be achieved, then a post-marketing study would be required to confirm clinical benefit. We considered a model in which drug-treated individuals are enrolled in a surveillance program and their survival is compared to that of historical controls. We estimate that, compared to randomized pre-approval studies, such a program could reduce the number of individuals required for 80% power at the *P*=0.05 threshold by 3-to 20-fold. Meanwhile, conditional approval of a first prion disease drug may alter key parameters such as diagnosis, recruitment, and genetic testing rates, the last of which alone could increase participant availability by more than 2-fold. Thus, while power for any trial depends upon how effective the drug is, there exists a range of assumptions under which a post-marketing study could be adequately powered. There may be various formats through which the Accelerated Approval requirement of a post-marketing study to confirm clinical benefit could be met.

Under some assumptions, a post-marketing study might last a decade or longer and would benefit from following all mutation carriers taking the drug. With creative and careful planning, we propose that these goals could be achieved. In one model, a post-marketing study might take the form of a surveillance program, in which treated patients are followed long-term, perhaps in collaboration with existing prion specialist clinics and surveillance centers worldwide. In such a model, drug costs would be reimbursed by payors, in contrast to a more traditional sponsor-funded pivotal trial. While this model would be a departure from the more conventional design of most post-marketing studies required for recent Accelerated Approval drugs^2^, precedents exist for regulatory innovation in this area. For example, FDA’s Risk Evaluation and Mitigation Strategies (REMS) program for drugs with serious safety concerns entails indefinite post-market enrollment and monitoring of treated patients^51^, and post-approval study requirements for medical devices often include registries or surveillance efforts and are not always industry-funded^52,53^.

Our study has several limitations. First, true age of onset distributions can only be obtained prospectively^54^, whereas our data are largely retrospective. We have included asymptomatic individuals with pathogenic *PRNP* variants where possible, but our ascertainment of them is certainly incomplete due to limited uptake of predictive testing^23^. This bias may tend to make our estimates of age of onset overly pessimistic^32,55^. To the extent that true age of onset is older, or total lifetime risk lower, than our data suggest, randomized preventive trials with a clinical endpoint would require even greater numbers of individuals, and thus further increase our caution around this study design. Second, although our dataset is, to our knowledge, the largest ever reported for genetic prion disease age of onset, our statistical power to detect genetic modifiers, which might aid in age of onset prediction, is still limited. Third, although we have attempted to select a reasonable set of assumptions for modeling clinical trials, we have by no means exhaustively sampled the set of possible trial designs and parameters. Fourth, powering a post-marketing study will require a good historical control dataset to compare to, and our dataset, which was collected mostly retrospectively, may or may not be adequate. We found no evidence that our dataset overestimates the hazards facing prospectively followed individuals, but this could be due to a lack of power in our analysis. Fifth, the ascertainment of genetic prion disease by prion surveillance centers may be biased towards rapidly progressive phenotypes, meaning that the prevalence of more slowly progressive forms might be underestimated.

Our findings highlight two priorities for the prion field. First, the discovery and characterization of biomarkers capable of serving as trial endpoints may be essential to enable near-term presymptomatic trials in genetic prion disease. Second, a post-approval surveillance mechanism for age of onset merits consideration as one option for confirmation of clinical benefit in the context of Accelerated Approval. The ability to access therapies that can prevent or delay prion disease, yet which are likely to be less effective or ineffective after symptom onset, could be greatly enhanced by success in these areas.

## Methods

### Literature annotation

We considered 69 reportedly pathogenic *PRNP* variants (Table S1) and reviewed primary literature to determine which had evidence of at least one family with at least three affected individuals in a pattern consistent with Mendelian segregation, or had a documented case with a *de novo* mutation. We identified 27 such variants, deemed likely high penetrance variants. The remainder were seen in isolated patient(s) with a negative or unknown family history, and/or have population allele frequencies inconsistent with high penetrance^6^. These variants will include both benign and low-risk variants. It is possible that some genuinely high penetrance variants may also lack literature evidence for high penetrance due to missing family history information or an unavailability of family member DNA to confirm *de novo* status, but this issue will only affect variants with very low case counts and thus will have minimal impact on the results reported here.

### Data collection

Age of onset data were gathered from nine study centers: the UK National Prion Clinic, the German Reference Center for TSEs, the Memory and Aging Center at University of California San Francisco, the Australian National CJD Registry, the reference center for CJD at University of Bologna, the DOXIFF study at the Mario Negri Institute, the Japanese national prion surveillance network, the French national reference center for CJD, and the Spanish National Center for Epidemiology. The data include both previously reported and newly identified families and individuals. Data were collected through clinical visits, reports to prion surveillance centers, and family histories, as previously described^32,56–61^. Age of onset was based on the earliest date of symptoms, determined by the patient or witnesses, that subsequently developed into prion disease. Data on the number of positive predictive genetic tests for *PRNP* mutations was provided by the National Prion Disease Pathology Surveillance Center for this study.

### Life tables and hazard curves

We tabulated, for each *PRNP* mutation and for each age from 1-100, the number of individuals alive at the beginning of the interval (lives; l), becoming sick or dying within the interval (deaths; d), or being censored – alive and well at last followup or dead of a different cause – within the interval (withdrawals; w). The raw hazard (q) was computed as onsets divided by the mean number of people observed over the interval: q = d/(l - w/2), and a smoothed hazard (q_smooth) was computed by passing a Gaussian filter (sd=3 years, maximum width=15 years) over the raw hazard. The proportion surviving for each interval (p) was 100% for the first year and was computed as (1-q) times the proportion surviving in the previous interval for every year thereafter. To compute the 95% confidence intervals on the smoothed hazard, we sampled each mutation’s data, with replacement, 1000 times, generated life tables for iteration, and then chose the 2.5^th^ and 97.5^th^ percentile of the hazards in the bootstrapped distributions at each age.

### Assumptions

To determine a reasonable assumption for withdrawal rate, we performed Google Scholar searches for preventive trials in neurology (N=2) or cardiology (N=6). The annual withdrawal rate was computed as w = 1 - exp(log(A)/t)), where A is the proportion of patients completing the trial at time t. Results are summarized in Table S3.

### Power calculation

The number of events (disease onsets, d) required was computed per Schoenfeld et al^30^ (equation 1). The number of patients required in order to observe that number of disease onsets was computed using an exponential model per Kohn et al^62^. Hazard in the placebo group was the baseline hazard specified in the text (4.6% for Table 2), and hazard for the drug group was the baseline hazard times the hazard ratio. The cumulative event rate in each group was computed as C = (h/(h+w)) * (1-exp(-(h + w)*t))), where h = hazard, w = withdrawal rate, and t = years of followup. The overall cumulative event rate C_tot_ was the average of the cumulative event rates for the two groups, weighted by proportion treated (in this case, 50/50). The number of randomized individuals required for d events to be observed was calculated as d /C_tot_. To account for ignoring the first *g* years of data, we reasoned that the cumulative rate of events usable in the final dataset would be C_usable_ = (h/(h+w)) * (1-exp(-(h + w)*t))) - (h/(h+w)) * (1-exp(-(h + w)*g))), which simplifies to C_usable_ = (h/(h+w)) * (exp(-(h + w)*g)- exp(-(h + w)*t))

### Simulations

Details of the simulations of randomized trials and historical control trials are in the Supplementary Discussion.

### Source code and data availability

Raw data cannot be made available due to identifiability concerns, but life tables have been included in supplement (Supplementary Life Tables). All analyses were conducted in the R programming language. Life tables and R source code are presented in a public GitHub repository at https://github.com/ericminikel/prnp_onset and are sufficient to reproduce most analyses and figures herein.

## Acknowledgments

This study was performed under ethical approval from the Partners Healthcare Institutional Research Board (2014P000226/MGH) and the Broad Institute’s Office of Research Subjects Protection (ORSP-2121 and NHSR-4190). We thank the patients and families who contributed their data to this research. EVM is supported by the National Institutes of Health (F31 AI122592), SV is supported by the National Science Foundation (GRFP 2015214731), and both are supported by an anonymous organization. SH and JPB are supported by Santé Publique France by the program “Investissement avenir” ANR-10-IAIHU-06. Work at the NHS National Prion Clinic was supported by the UK National Institute of Health Research’s Biomedical Research Centre at University College London Hospitals NHS Foundation Trust. Japanese prion surveillance is funded by a grant-in-aid from the Research Committee of Prion Disease and Slow Virus Infection, the Ministry of Health, Labour and Welfare of Japan, and from the Research Committee of Surveillance and Infection Control of Prion Disease, the Ministry of Health, Labour and Welfare of Japan. SC is supported in part by an NHMRC Practitioner Fellowship (#APP1105784) and the Australian National Creutzfeldt-Jakob Disease Registry is funded by the Commonwealth Department of Health. Data collection in Germany was supported by a grant number 1369-341 by Bundesministerium für Gesundheit through Robert Koch Institute. Work in Italy was partially supported by Fondazione Telethon (GGP10208). We thank Mr. Javier Almazan and Dr. Maria Ruiz, CNE, Madrid, for help with data extraction at the Spanish CJD Registry. We thank Eric S. Lander, Michael J. Donovan, and Adam Bliss for discussions.

## Supplementary Discussion

### Estimation of number of individuals available for trials

It is possible to estimate the true number of high penetrance *PRNP* mutation carriers based on disease prevalence. Using data from recent case series, 1,176 prion disease cases harbored a *PRNP* variant classified here as highly penetrant, out of 10,460 sequenced cases or 16,025 total cases^6^. Thus, 7 - 11% of prion disease cases have a high penetrance *PRNP* variant. Prion disease is responsible for ∼1 in 5,000 deaths^6^, suggesting that ∼1 in 45,000 to 71,000 deaths are due to a high penetrance *PRNP* variant. The carrier rate among the living population will be somewhat lower because these variants reduce life expectancy, but it is reasonable to suppose that ∼1 in 100,000 people harbors a high penetrance *PRNP* variant. This is in line with recent population control data, where out of 138,632 individuals in the gnomAD database as of December 2017 (http://gnomad.broadinstitute.org/)^63^, there is one individual with the E200K mutation and no others with any variant classified here as high penetrance. Similarly, out of ∼531,575 individuals genotyped by 23andMe, between 1 and 5 harbored one of four well-known high penetrance variants (P102L, A117V, D178N, and E200K) and between 1 and 5 harbored one of an additional set of variants which includes three classified here as high penetrance (P105L, T183A, and F198S). If the true carrier rate is 1 in 100,000, then there may exist 3,000 people in the United States with high penetrance *PRNP* variants. However, this figure greatly overestimates the number of people available for trials, as most of these individuals have not undergone predictive testing. Indeed, many are likely not even aware that they are at risk, perhaps because a family history is absent or a family member was not diagnosed correctly.

The National Prion Disease Pathology Surveillance Center in Cleveland, Ohio, as the sole provider of *PRNP* gene testing in the U.S., has exhaustive ascertainment of individuals who have chosen predictive testing for genetic prion disease in this country. In the period from 1996 through January 2017, it provided *N*=221 positive predictive test results, for any *PRNP* variant, to individuals who are not known to have developed disease as of 2017. Privacy concerns prevent publication of a breakdown of this number by age and specific *PRNP* mutation, but estimates can be made based on other cohorts. Among U.S. symptomatic prion disease cases with a rare *PRNP* variant, 75% (271/362) of individuals had a mutation classified here as high penetrance (Table S1), and in the reported U.K. predictive testing cohort^23^, 36% (37/104) of individuals who chose predictive testing were age 40 or older. Thus, a conservative estimate that there are only 221× 75%×36% = ∼60 individuals alive in the United States today who meet the criteria we use in our power calculations.

The above estimate is conservative in that it reflects individuals who currently know their genetic status. In the U.K. predictive testing cohort, only 23% of individuals at 50/50 risk chose predictive testing^23^, similar to reported figures in Huntington’s disease, another incurable neurological disease (see refs in ^23^). In contrast, 60% of individuals at risk for *BRCA1* or *BRCA2* mutations, for which preventive measures are available, chose predictive testing^37^. Thus, it is possible to imagine a 60%/23% = ∼2.6X increase in the uptake of predictive testing if a preventive therapy for prion disease were available. Thus, a more generous estimate of the number of individuals age ≥40 available in the U.S. is 60×2.6 = 156. Such an estimate is probably more realistic when considering an approved prevention measure (as in post-marketing studies) than when considering an experimental drug entering randomized pre-approval trials (see below).

Although we contemplated worldwide trials with multiple international sites, we did not have adequate data to estimate the number of genetically tested presymptomatic individuals worldwide. The NHS National Prion Clinic in the U.K. has seen 72 presymptomatic individuals with *PRNP* mutations since 1990, and the French surveillance center in Paris has delivered 18 positive *PRNP* predictive test results since 2004, but the other centers involved in this report did not have comprehensive data on predictive testing in their respective countries analogous to that available for the U.S. We also note that a large number of E200K mutation carriers are suspected to exist in Slovakia and Israel due to founder mutations, although fewer than 100 carriers appear to have been identified in each country to date^64,65^.

We also considered estimates based on the incidence of genetic prion disease. U.S. prion surveillance reported 271 individuals dying of prion disease with high penetrance mutations, suggesting that at least a comparable number of carriers in the U.S. are currently healthy and will have onset with a correct diagnosis within the next 15 years. The comparable figure including Europe, Australia, and Japan is 1,176. These last two figures are still lower than the true number of carriers in existence due to underdiagnosis, yet they overestimate the number of individuals actually reachable for trials because they ignore the question of how many individuals would choose predictive testing, and they include individuals who would be difficult to ascertain prospectively because they lack a known family history of prion disease, either due to *de novo* mutations, incorrect or incomplete information about previous family illnesses, or <100% penetrance.

For all of the above estimates, an important caveat is that the number of individuals successfully recruited, screened, and enrolled for a trial will be only a fraction of the number who meet the most basic enrollment criteria such as genetic status and age. Willingness, geography, and various exclusion criteria will dramatically lower the number actually enrolled.

Finally, it is worth noting that all of our calculations and assumptions are based upon the present moment, when there exists no drug for prion disease. It is likely that approval of a first prion disease drug would increase the number of patients available for future trials. A drug could improve diagnosis rates, as prion disease is not currently prioritized in the differential diagnosis of rapidly progressive dementia due to its being untreatable^66^. The U.S. observes an incidence of ∼1 prion disease case per million population per year, but up to twice that incidence has been observed in countries with more intense surveillance systems^29^. Because many prion disease patients die undiagnosed, their relatives may never learn that they are at risk for a *PRNP* mutation. A drug might also increase the uptake of predictive genetic testing among those who do learn that they are at risk. The 23% uptake observed for prion disease^23^ is consistent with other currently “medically inactionable” indications such as Huntington’s disease^67^, while as noted in the main text, “actionable” indications such as *BRCA1/2* mutations appear to have much higher uptake^37^. Finally, the existence of a drug may promote general awareness of the disease and improve the infrastructure for surveillance, registries, and patient ascertainment.

### Simulation of power for randomized preventive trials with a clinical endpoint

Individuals were assigned one of the three *PRNP* mutations and a starting age distributed between 40 and 80, weighted by mutation prevalence and by the proportion of individuals surviving at each age. As above, we assigned half of individuals to drug and half to placebo, and assumed a w=15.2% annual withdrawal rate, a *P*=0.05 statistical threshold, and a 5-year trial duration with a 1-year “run-in” period. For each year of the trial, each individual withdraws with probability w, becomes sick with a probability corresponding to the hazard function for their particular *PRNP* mutation and age at the time, multiplied by the simulated hazard ratio if drug treated, or else continues on in the trial. At the end of each simulated trial, we analyzed the censored trial data to determine a *P* value. For non-stratified simulations, drug/placebo status was assigned without regard to mutation, and survival status was regressed on drug/placebo status alone using a log-rank test, with the overall *P* value as the readout. For stratified trials, drug/placebo status was assigned 50/50 within each mutation, and mutation was included as a covariate in a Cox proportional hazards regression, with the *P* value for the “drug” parameter as the readout.

We then compared this model to the power calculation results by taking the calculated required numbers of individuals for 80% power (Table 2) and then running the simulation (500 iterations) to determine the power for this number of individuals. The results (Table S6) show overall good agreement between the power calculation and the simulation — for most scenarios tested, the power is indeed close to 80%, with or without stratification. Stratification actually reduces statistical power for the conditions with low *N* and low hazard ratios. Under such conditions, it is a common occurrence that there may be zero disease onsets either in one randomized group (usually the drug-treated group) or in one mutation, resulting in an infinite regression coefficient or beta in the Cox model. Thus, the regression never converges, and the simulated trial results turn out statistically non-significant.

### Codon 129 effects on age of onset and disease duration

To determine whether codon 129 affects age of onset for the three most prevalent mutations considered here, we used a log-rank model based on codon 129 diplotype (phased genotype) where available (Table S7). In this model, only D178N showed clear evidence for genetic modification of age of onset and disease duration, with *P* values significant after multiple testing correction. To determine the nature of this genetic modification, we plotted survival curves by codon 129 diplotype and, because phase was unknown for many codon 129 heterozygous individuals, we also considered phaseless genotypes. In pairwise tests for D178N, M/M was not significantly different from M/V (nominal *P* = 0.14) nor from V/M (nominal *P* = 0.69), and in the phaseless survival curve, MV was overall similar to MM (Figure S4D). These results suggest that the significant codon 129 effect on D178N age of onset is most likely driven primarily by a younger age of onset in V/V individuals compared to other diplotypes. Despite the strong statistical significance of this difference, the small number of D178N-129VV individuals means that codon 129 does not add any explanatory power for age of onset in the dataset as a whole. As noted in the main text, mutation alone explains limited variance in age of onset (adjusted R^2 = 0.15, *P* = 1.3e-33). Adding *cis* and *trans* codon 129 to this model decreases the variance explained (adjusted R^2 = 0.14, *P* = 3.6e-18).

We also investigated in further detail previously reported associations. For disease duration, D178N M/M and V/V were significantly more rapid than either heterozygous diplotype, consistent with previous reports. Although codon 129 diplotype did not have a significant effect on E200K disease duration overall (nominal *P* = 0.10), a phaseless genotypic model was suggestive (nominal *P* = 0.031), with MV heterozygotes appearing to have a slightly longer disease duration than MM homozygotes, a direction of effect consistent with previous reports^5,32^. Whereas P102L age of onset was reported to be higher for M/V than M/M individuals^36^, here we find no evidence for this and, ignoring phase, the non-significant trend is towards younger onset in MV than MM individuals (nominal *P* = 0.056).

### Potential age of onset confounders

Because our data were gathered from a variety of study centers using a variety of methodologies, we asked whether any confounders might affect age of onset (Table S7). There was no difference in age of onset between directly and indirectly ascertained individuals (*P* = 0.78). Age of onset was correlated with year of birth after controlling for mutation (*P* < 1e-48), which is a previously reported artifact caused by our relatively limited ability to ascertain individuals whose onset has not yet arrived (though we ascertain some of them through predictive testing) or whose onset occurred before genetic diagnosis of prion disease was possible (though we ascertain some of them through family histories)^32^. This correlation does not affect estimation of overall age of onset distributions. Age of onset appeared to differ slightly among the nine contributing study centers after controlling for mutation, although it was not significant after multiple testing correction (nominal *P* = 0.012, Bonferroni *P* = 0.26, two-way ANOVA), and it only marginally increased variance explained (adjusted R^2 = 0.16) compared to mutation alone (adjusted R^2 = 0.15, see above). Year of onset showed evidence of positive correlation with age of onset after controlling for study center and mutation (nominal *P* = 0.00032, Bonferroni *P* = 0.008, linear regression), although the effect size was small (+0.12 years of age per calendar year, or in other words, cases in 2010 have on average an age of onset 1.2 years older than cases in 2000) and, again, the impact on variance explained was minimal (adjusted R^2 = 0.18). This slight positive correlation might be due to improved ascertainment of older-onset cases as prion surveillance strengthens over time.

### Justification for trial duration assumptions

In the main text, we argued that a longer trial duration could be considered for a post-marketing study because it would run concurrently with, rather than reducing, the drug’s effective market exclusivity period (the period before generic equivalents can be approved). In the U.S., new drugs may be protected by patent exclusivity granted by the Patent and Trademark Office and/or by market exclusivity measures granted by FDA; these exclusivity periods are not additive. Patents last 20 years beginning from their filing, which is usually during the preclinical development phase. The 1984 Hatch-Waxman Act allows sponsors to recover up to 5 years of additional exclusivity, not to exceed a total of 14 years of market exclusivity, to make up for time the drug spends in FDA review^68^. FDA can offer varying periods of market exclusivity depending upon the indication and treatment modality, including 12 years for new biologics^69^ and 7 years for rare disease drugs granted Orphan Drug designation^70^. In practice, new drugs receive on average about 12 years of effective market exclusivity^71,72^. The vast majority of pivotal trials supporting new drug approvals last less than one year^73^. While there are rare examples of 5-year trials^47^, a 10-or 15-year prevention trial would exhaust most or all of a drug’s effective market exclusivity period. In contrast, as noted in the Discussion, there do exist precedents for very long-term surveillance of patients receiving a drug after approval.

### Historical control trial simulation

As for the simulation of randomized trials, individuals were assigned one of the three *PRNP* mutations and a starting age distributed between 40 and 80, weighted by mutation prevalence and by the proportion of individuals surviving at each age. Again, we assumed a w=15.2% annual withdrawal rate (Table S5), a *P*=0.05 statistical threshold, and a 1-year “run-in” period where disease events are ignored. Distinct from the randomized trial simulation, here all simulated individuals are treated with the drug. For each year of the trial, each individual withdraws with probability w, becomes sick with a probability corresponding to the hazard function for their particular *PRNP* mutation and age at the time, multiplied by the simulated hazard ratio, or else continues on in the trial. At the end of each simulated trial, the censored trial data on treated individuals are compared to our original dataset as historical controls (Supplementary Life Tables). To determine a *P* value we used a Cox proportional hazards counting model accounting for different left-truncation times^74^: for untreated individuals in the original dataset, we assumed age 0 as a start time, while for treated individuals, we assumed left truncation at the age at trial enrollment, plus one year to account for the “run-in” year.

While we cannot currently rule out the possibility that our dataset is biased relative to the *true* hazards facing mutation carriers in real life (see main text Discussion), we sought to confirm that our simulation method is not itself biased. We reasoned that if our simulation was unbiased, then for a drug with hazard ratio equal to 1 (a completely ineffective drug), even long trials with large numbers of individuals should have power equal to alpha, by the definition that alpha is the false positive rate when the null hypothesis (no efficacy) is true. We therefore ran 1000 iterations of a simulation with a hazard ratio of 1 and 1000 individuals followed for 20 years. We observed a significant result at P < 0.05 in only 5.5% of iterations, consistent with the expected 5%.

In contrast to the result for randomized trials (see discussion above and Table S6), we found that stratification by mutation in the analysis of historical control trial simulation did just slightly increase statistical power. For example, with *N*=156 individuals followed for 15 years with a hazard ratio of 0.5, power was 90.6% (906/1000 iterations) without stratification and 94.1% (941/1000 iterations) with stratification. This difference from the randomized trial simulation may be a property of the Cox counting model, combined with the fact that our historical comparison dataset has *N*=1,000 individuals, and we considered follow-up periods of up to 15 years, meaning that the dataset was large enough for the small explanatory power of different *PRNP* mutations to matter. Nevertheless, for consistency with the methods used for the randomized trial simulations, we chose not to stratify in the simulations used for Table 3 and Figure S6.

We performed power calculations for post-marketing studies using historical controls under a range of assumptions in addition to those explored in Table 3 in the main text. In one set of experiments, we considered the effects of varying the length of the follow-up period. For a hazard ratio of 0.5, 80% power could be achieved within 9 years for *N*=156 participants, but is never achieved for *N*=60 participants (Figure S6A). This is because statistical power eventually plateaus for lack of participants: our assumption of a 15.2% withdrawal rate compounded annually means that after 10 years, only 19% of the original participants remain in the trial. If the set of drug recipients followed in a post-marketing study were fixed shortly after approval, then this is a realistic concern. If, on the other hand, study design allows new individuals who are prescribed the drug to be added to the monitored cohort continually, the number of individuals in the trial could stay constant or even grow. To simulate this possibility, we also considered a zero withdrawal rate scenario. Under this assumption, even with *N*=60 individuals, 80% power is achieved in 10 years (Figure S6A).

In another set of experiments, we compared the power for post-marketing studies with historical controls, with or without modeling withdrawal, in comparison to pre-approval randomized trials, for a range of hazard ratios (Figure S6B). For the same hazard ratio and level of statistical power, post-marketing trials generally required only about one fifth as many individuals, and if withdrawal is set to zero, simulating continuous enrollment, only one twentieth as many, as pre-approval randomized trials.

Certainly, a post-marketing study is not a panacea, and under certain assumptions even this trial design is not well-powered: for instance, for a drug of marginal efficacy (hazard ratio 0.9, delaying onset by ∼1 year) even a 15-year trial with no withdrawal could not achieve 80% power with 1,000 participants. But, under a range of moderate assumptions, a post-marketing study is more feasible than randomized pre-approval trials with a clinical endpoint.

## Supplementary Tables

**Table S1.**
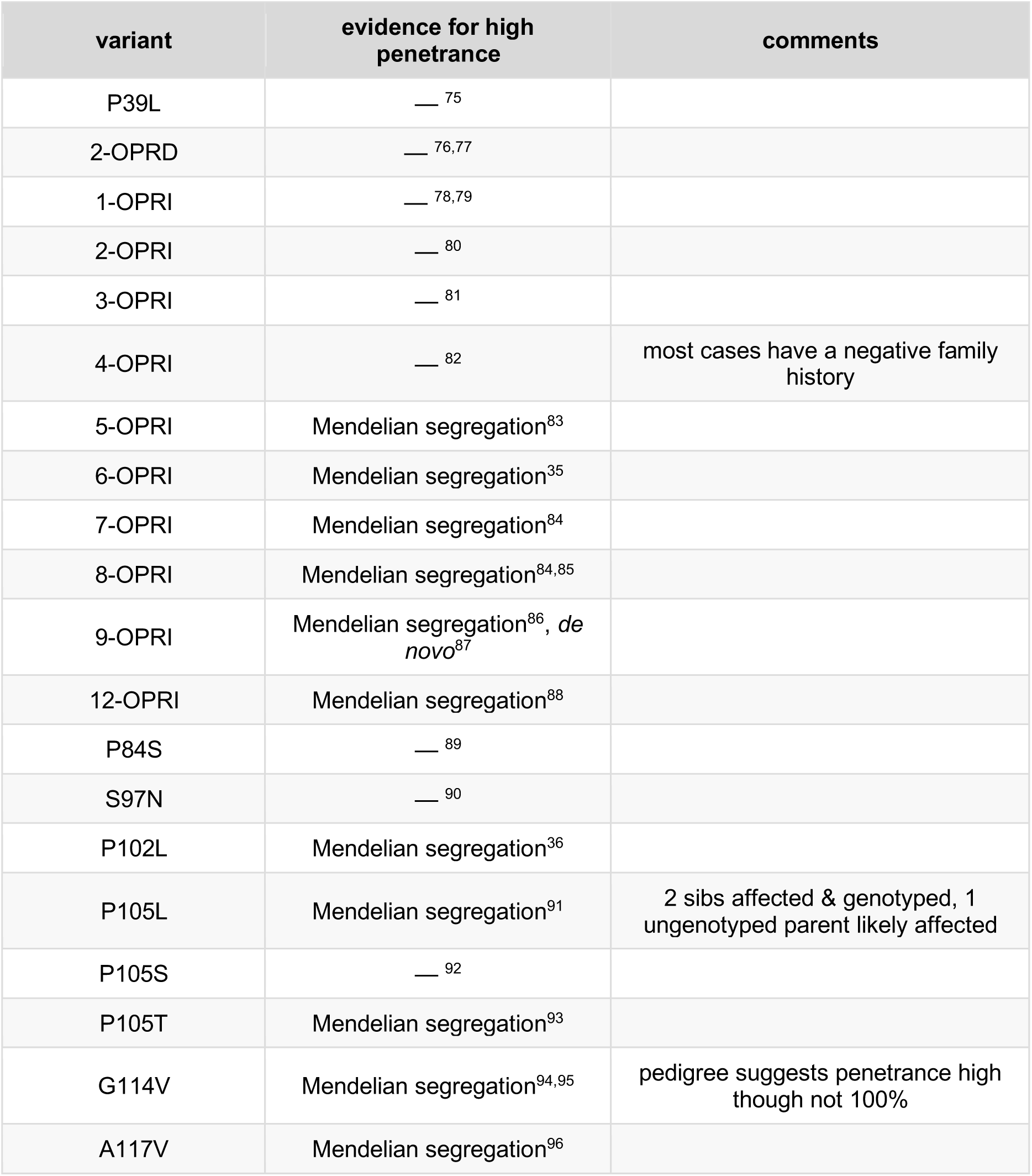
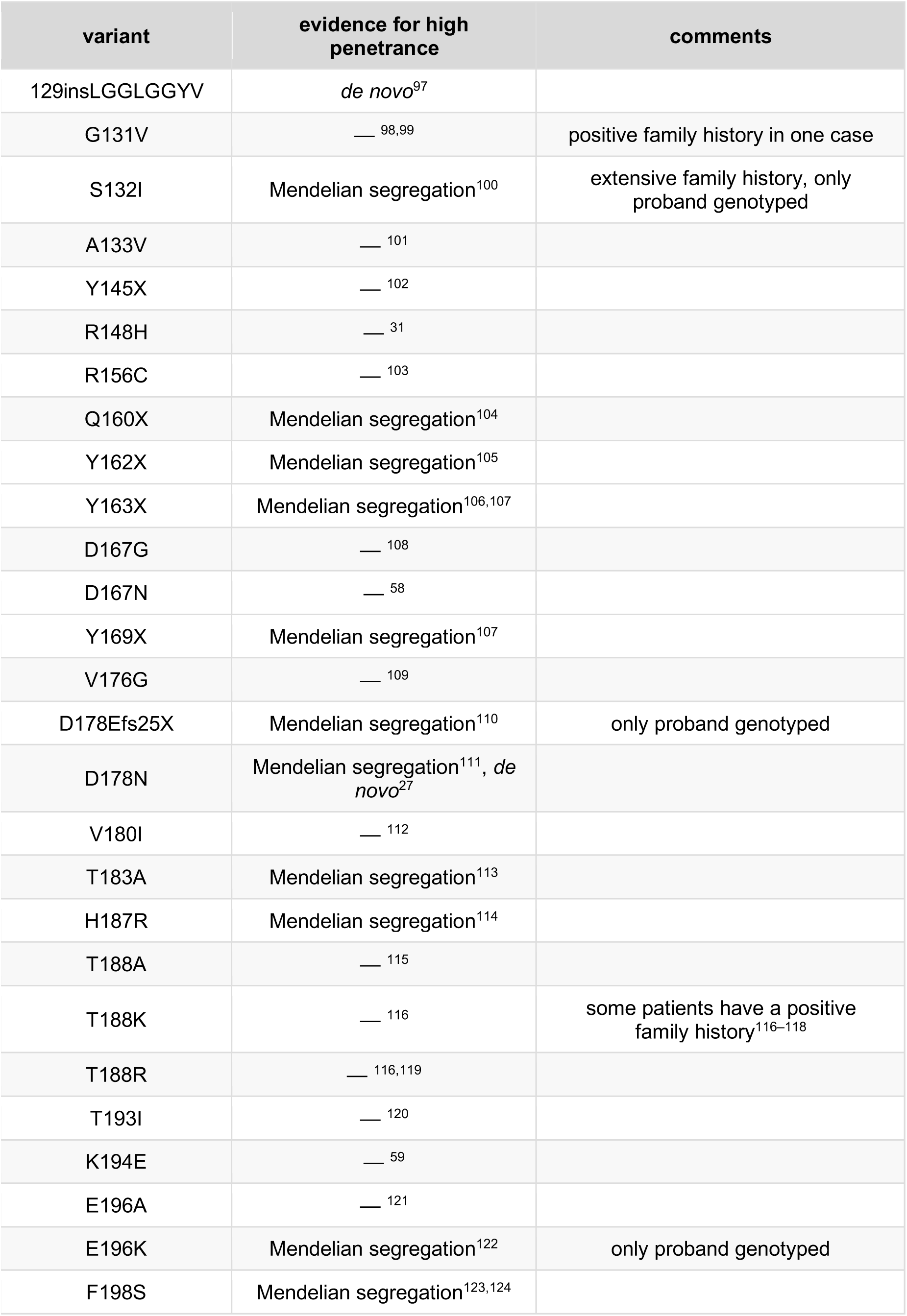
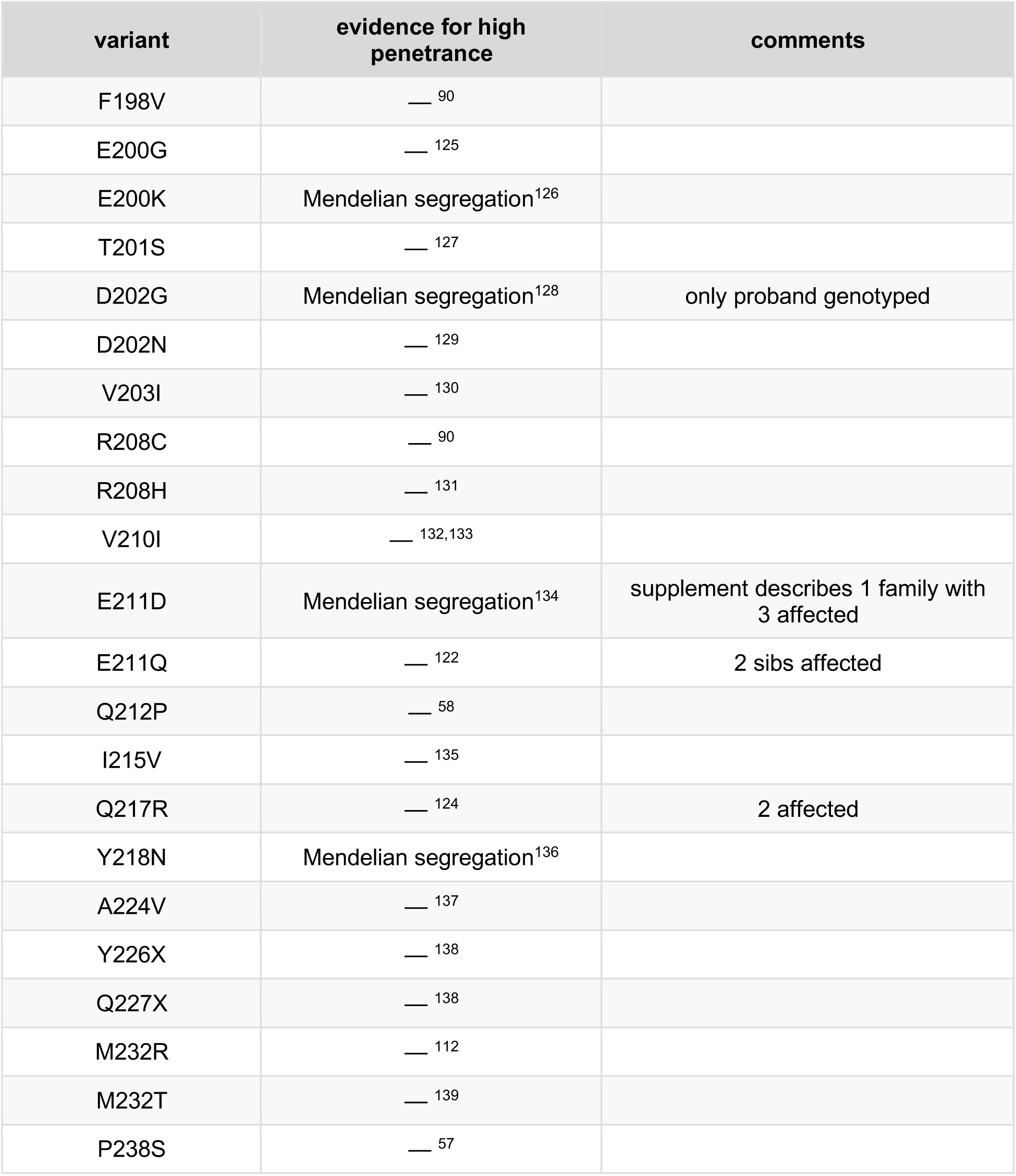
Literature review to identify probable high penetrance variants. “Mendelian segregation” indicates the presence of at least one family with at least three affected individuals in a pattern consistent with Mendelian segregation. “De novo” indicates a case with a confirmed de novo mutation. — indicates neither of these criteria was present.

| variant | evidence for high penetrance | comments |
| --- | --- | --- |
| P39L | — <sup>75</sup> |  |
| 2-OPRD | — <sup>76,77</sup> |  |
| 1-OPRI | — <sup>78,79</sup> |  |
| 2-OPRI | — <sup>80</sup> |  |
| 3-OPRI | — <sup>81</sup> |  |
| 4-OPRI | — <sup>82</sup> | most cases have a negative family history |
| 5-OPRI | Mendelian segregation<sup>83</sup> |  |
| 6-OPRI | Mendelian segregation<sup>35</sup> |  |
| 7-OPRI | Mendelian segregation<sup>84</sup> |  |
| 8-OPRI | Mendelian segregation<sup>84,85</sup> |  |
| 9-OPRI | Mendelian segregation<sup>86</sup>, *de novo*<sup>87</sup> |  |
| 12-OPRI | Mendelian segregation<sup>88</sup> |  |
| P84S | — <sup>89</sup> |  |
| S97N | — <sup>90</sup> |  |
| P102L | Mendelian segregation<sup>36</sup> |  |
| P105L | Mendelian segregation<sup>91</sup> | 2 sibs affected & genotyped, 1 ungenotyped parent likely affected |
| P105S | — <sup>92</sup> |  |
| P105T | Mendelian segregation<sup>93</sup> |  |
| G114V | Mendelian segregation<sup>94,95</sup> | pedigree suggests penetrance high though not 100% |
| A117V | Mendelian segregation<sup>96</sup> |  |
| 129insLGGLGGYV | *de novo*<sup>97</sup> |  |
| G131V | — <sup>98,99</sup> | positive family history in one case |
| S132I | Mendelian segregation<sup>100</sup> | extensive family history, only proband genotyped |
| A133V | — <sup>101</sup> |  |
| Y145X | — <sup>102</sup> |  |
| R148H | — <sup>31</sup> |  |
| R156C | — <sup>103</sup> |  |
| Q160X | Mendelian segregation<sup>104</sup> |  |
| Y162X | Mendelian segregation<sup>105</sup> |  |
| Y163X | Mendelian segregation<sup>106,107</sup> |  |
| D167G | — <sup>108</sup> |  |
| D167N | — <sup>58</sup> |  |
| Y169X | Mendelian segregation<sup>107</sup> |  |
| V176G | — <sup>109</sup> |  |
| D178Efs25X | Mendelian segregation<sup>110</sup> | only proband genotyped |
| D178N | Mendelian segregation<sup>111</sup>, *de novo*<sup>27</sup> |  |
| V180I | — <sup>112</sup> |  |
| T183A | Mendelian segregation<sup>113</sup> |  |
| H187R | Mendelian segregation<sup>114</sup> |  |
| T188A | — <sup>115</sup> |  |
| T188K | — <sup>116</sup> | some patients have a positive family history<sup>116–118</sup> |
| T188R | — <sup>116,119</sup> |  |
| T193I | — <sup>120</sup> |  |
| K194E | — <sup>59</sup> |  |
| E196A | — <sup>121</sup> |  |
| E196K | Mendelian segregation<sup>122</sup> | only proband genotyped |
| F198S | Mendelian segregation<sup>123,124</sup> |  |
| F198V | — 90 |  |
| E200G | — 125 |  |
| E200K | Mendelian segregation<sup>126</sup> |  |
| T201S | — 127 |  |
| D202G | Mendelian segregation<sup>128</sup> | only proband genotyped |
| D202N | — 129 |  |
| V203I | — 130 |  |
| R208C | — 90 |  |
| R208H | — 131 |  |
| V210I | — 132,133 |  |
| E211D | Mendelian segregation<sup>134</sup> | supplement describes 1 family with 3 affected |
| E211Q | — 122 | 2 sibs affected |
| Q212P | — 58 |  |
| I215V | — 135 |  |
| Q217R | — 124 | 2 affected |
| Y218N | Mendelian segregation<sup>136</sup> |  |
| A224V | — 137 |  |
| Y226X | — 138 |  |
| Q227X | — 138 |  |
| M232R | — 112 |  |
| M232T | — 139 |  |
| P238S | — 57 |  |

**Table S2.** Descriptive statistics regarding sources of age of onset data.

| **study center** | **N** |
| --- | --- |
| Japanese national prion surveillance network (Shimotsuke & Kanazawa, Japan) | 215 |
| MRC Prion Unit (London, U.K.) | 211 |
| French national reference center for CJD (Paris, France) | 168 |
| UCSF Memory and Aging Center (San Francisco, U.S.) | 147 |
| Spanish National Center for Epidemiology (Madrid, Spain) | 114 |
| German Reference Center for TSEs (Göttingen, Germany) | 101 |
| DOXIFF study at the Mario Negri Institute (Milan, Italy) | 65 |
| Reference Center for CJD at University of Bologna (Bologna, Italy) | 49 |
| Australian National CJD Registry (Melbourne, Australia) | 24 |
| *total* | *1094* |
| **method of ascertainment** | **N** |
| direct (clinical visit, autopsy, or surveillance report) | 843 |
| indirect (family history) | 251 |
| *total* | *1094* |
| **vital status** | **N** |
| censored — died due to intercurrent illness without developing prion disease | 4 |
| censored — alive and well at last follow-up | 101 |
| symptomatic with prion disease at last follow-up | 81 |
| died of prion disease | 908 |
| *total* | *1094* |

**Table S3.** Age of onset statistics on supplementary variants.

| **mutation** | **without censored data** |  | **survival curve including censored data** |  |  |
| --- | --- | --- | --- | --- | --- |
|  | **mean ± sd** | **N** | **median (IQR)** | **range** | **N** |
| 5-OPRI | 46.8 ± 6.0 | 14 | 49 (44 - 53) | 34 - 56 | 18 |
| 6-OPRI | 35.1 ± 5.8 | 31 | 35 (32 - 39) | 23 - 47 | 34 |
| P105L | 46.5 ± 8.5 | 13 | 47 (40 - 51) | 31 - 61 | 13 |
| A117V | 41.2 ± 7.8 | 26 | 41 (37 - 45) | 25 - 58 | 28 |

**Table S4.** Withdrawal rates in preventive clinical trials. w, annual withdrawal rate. CHD, coronary heart disease. NSAID, non-steroidal anti-inflammatory drug. See Methods for details.

| category | trial | description | w |
| --- | --- | --- | --- |
| cardiology | WOSCOPS<sup>140</sup> | pravastatin for CHD | 6.9% |
| cardiology | AFCAPS/TexCAPS<sup>141</sup> | lovastatin for CHD | 7.1% |
| cardiology | OSLER<sup>142</sup> | evolocumab for CHD | 9.0% |
| cardiology | JUPITER<sup>143</sup> | rosuvastatin for CHD | 14.1% |
| neurology | ADAPT<sup>144</sup> | NSAIDs for Alzheimer's | 16.3% |
| cardiology | ODYSSEY LONG TERM<sup>145</sup> | alirocumab for CHD | 19.0% |
| cardiology | NCT00607373<sup>146</sup> | mipomersen for homozygous *LDLR* hypercholesterolemia | 22.1% |
| neurology | PRECREST<sup>147</sup> | creatine for Huntington's disease | 54.9% |

**Table S5.** Power calculations under alternative assumptions. Each block of this table is equivalent to Table 3 but with different assumptions as indicated (except where stated, other assumptions are identical to those in Table 2). A) Best case scenario: overall average hazard is 4.8% (the higher figure including the less common mutations shown in Table S2 and Figure S1), the withdrawal rate is 6.9% per year (the lowest rate in any of the trials we reviewed, see Table S4), and there is no run-in period — the drug is effective immediately and so disease onsets within the 1st year of the trial are included. B) Worst case scenario: overall average hazard is only 3.5% — one quarter lower than calculated in this manuscript, because our data are biased due to under-incluson of asymptomatic individuals, and/or because predominantly younger people enroll in a trial — and the withdrawal rate is 54.9% per year (the highest rate in any trial we reviewed, see Table S4). C) Targeted trial scenario: only the mutations with higher hazards — 5-OPRI, 6-OPRI, P105L, and A117V — are targeted for recruitment, resulting in a higher baseline hazard of 5.2%. Although the enrollment requirements for this scenario are lower than in Table 2, these mutations are also approximately one order of magnitude rarer^6^, making achievement of these enrollment numbers yet more unlikely. D) Long follow-up scenario: trial duration is 15 years. This reduces the required numbers somewhat, but this benefit is limited by the withdrawal rate, which means that few individuals are still enrolled after 15 years. E) Zero withdrawal scenario: withdrawal rate is set to zero.

| alternate scenario | hazard ratio | years of life added | onsets required | participants required |
| --- | --- | --- | --- | --- |
| A<br>(best case) | 0.1 | undefined* | 6 | 59 |
|  | 0.2 | 21 | 12 | 110 |
|  | 0.3 | 13 | 22 | 182 |
|  | 0.4 | 9 | 37 | 291 |
|  | 0.5 | 7 | 65 | 475 |
|  | 0.6 | 5 | 120 | 821 |
|  | 0.7 | 3 | 247 | 1,589 |
|  | 0.8 | 2 | 631 | 3,850 |
|  | 0.9 | 1 | 2,828 | 16,434 |
| B<br>(worst case) | 0.1 | undefined* | 6 | 357 |
|  | 0.2 | 21 | 12 | 666 |
|  | 0.3 | 14 | 22 | 1,097 |
|  | 0.4 | 10 | 37 | 1,757 |
|  | 0.5 | 7 | 65 | 2,866 |
|  | 0.6 | 5 | 120 | 4,955 |
|  | 0.7 | 4 | 247 | 9,586 |
|  | 0.8 | 2 | 631 | 23,196 |
|  | 0.9 | 1 | 2,828 | 98,897 |
| C<br>(targeted trial) | 0.1 | undefined* | 6 | 92 |
|  | 0.2 | undefined* | 12 | 171 |
|  | 0.3 | 10 | 22 | 280 |
|  | 0.4 | 7 | 37 | 449 |
|  | 0.5 | 5 | 65 | 732 |
|  | 0.6 | 4 | 120 | 1,267 |
|  | 0.7 | 3 | 247 | 2,457 |
|  | 0.8 | 2 | 631 | 5,958 |
|  | 0.9 | 1 | 2,828 | 25,471 |
| D<br>(long follow-up) | 0.1 | undefined* | 6 | 59 |
|  | 0.2 | 21 | 12 | 108 |
|  | 0.3 | 14 | 22 | 178 |
|  | 0.4 | 10 | 37 | 285 |
|  | 0.5 | 7 | 65 | 465 |
|  | 0.6 | 5 | 120 | 806 |
|  | 0.7 | 4 | 247 | 1,568 |
|  | 0.8 | 2 | 631 | 3,816 |
|  | 0.9 | 1 | 2,828 | 16,384 |
| E<br>(zero withdrawal) | 0.1 | undefined* | 6 | 66 |
|  | 0.2 | 21 | 12 | 123 |
|  | 0.3 | 14 | 22 | 202 |
|  | 0.4 | 10 | 37 | 323 |
|  | 0.5 | 7 | 65 | 527 |
|  | 0.6 | 5 | 120 | 912 |
|  | 0.7 | 4 | 247 | 1,767 |
|  | 0.8 | 2 | 631 | 4,285 |
|  | 0.9 | 1 | 2,828 | 18,314 |

**Table S6.** Comparison of power calculation and simulation results. The first four columns are reproduced from Table 2 for ease of comparison. The number of participants required was calculated to yield 80% power; the final two columns show the power for this number of participants, at P=0.05, indicated by simulation. See Supplementary Discussion above for details of the method.

| hazard ratio | years of life added | onsets required | participants required | calculated power | simulated power without stratification | simulated power with stratification |
| --- | --- | --- | --- | --- | --- | --- |
| 0.1 | undefined* | 6 | 101 | 80.0% | 62.2% | 35.0% |
| 0.2 | 21 | 12 | 189 | 80.0% | 71.8% | 69.2% |
| 0.3 | 14 | 22 | 311 | 80.0% | 78.6% | 76.6% |
| 0.4 | 10 | 37 | 498 | 80.0% | 80.0% | 81.2% |
| 0.5 | 7 | 65 | 813 | 80.0% | 80.0% | 81.8% |
| 0.6 | 5 | 120 | 1406 | 80.0% | 83.2% | 79.8% |
| 0.7 | 4 | 247 | 2,724 | 80.0% | 80.4% | 80.8% |
| 0.8 | 2 | 631 | 6,602 | 80.0% | 81.4% | 77.6% |
| 0.9 | 1 | 2,828 | 28,204 | 80.0% | 80.6% | 78.0% |

**Table S7.** Tests for modifiers and confounders of age of onset. All p-values are two-tailed. As explained in Supplementary Discussion, diplotypes (phased genotypes) are indicated with a slash (cis/trans to the mutation) while unphased genotypes have no slash. We were unable to obtain phase data for many 129MV individuals, so the genotypic tests represent not only a different grouping of data but also include more data points than the corresponding diplotypic tests. Thus, we considered them as independent tests for the purposes of multiple testing correction. p (raw) indicates the raw p value; p (bc) is Bonferroni-corrected for 22 tests. For parent-child comparisons n is the number of pairs. For linear regressions, child year of birth was included in the model as a covariate. The prior column represents the prior expectation of whether there would be a significant difference in each test based on previous reports in the literature.

| variable | mutation | comparison | n | test | *P* (raw) | *P* (bc) | prior evidence |
| --- | --- | --- | --- | --- | --- | --- | --- |
| onset | P102L | M/M vs. M/V vs. V/V | 125 vs. 13 vs. 1 | log-rank | 0.18 | 1 | mixed<sup>21,35,36</sup> |
| onset | P102L | MM vs. MV vs. VV | 125 vs. 32 vs. 1 | log-rank | 0.057 | 1 |  |
| onset | D178N | M/M vs. M/V vs. V/M vs. V/V | 133 vs. 18 vs. 9 vs. 13 | log-rank | 0.000062 | 0.0014 | none<sup>21,28,34,148</sup> |
| onset | D178N | MM vs. MV vs. VV | 133 vs. 58 vs. 13 | log-rank | 0.000018 | 0.00040 |  |
| onset | E200K | M/M vs. M/V vs. V/M vs. V/V | 286 vs. 33 vs. 10 vs. 5 | log-rank | 0.13 | 1 | none for *trans* allele<sup>21,32,34,149</sup>, suggestive for *cis* allele<sup>32</sup> |
| onset | E200K | MM vs. MV vs. VV | 288 vs. 92 vs. 5 | log-rank | 0.30 | 1 |  |
| duration | P102L | M/M vs. M/V | 89 vs. 8 | log-rank | 0.55 | 1 | none<sup>36</sup> |
| duration | P102L | MM vs. MV | 89 vs. 21 | log-rank | 0.93 | 1 |  |
| duration | D178N | M/M vs. M/V vs. V/M vs. V/V | 62 vs. 13 vs. 8 vs. 10 | log-rank | 0.000081 | 0.0018 | yes<sup>28</sup> |
| duration | D178N | MM vs. MV vs. VV | 62 vs. 33 vs. 10 | log-rank | 0.00000010 | 0.0000022 |  |
| duration | E200K | M/M vs. M/V vs. V/M vs. V/V | 208 vs. 21 vs. 6 vs. 5 | log-rank | 0.10 | 1 | yes<sup>5,32,64</sup> |
| duration | E200K | MM vs. MV vs. VV | 210 vs. 50 vs. 5 | log-rank | 0.031 | 0.68 |  |
| onset | P102L | parent vs. child | 32 | linear regression | 0.44 | 1 | suggestive<sup>150</sup> |
| onset | D178N | parent vs. child | 15 | linear regression | 0.12 | 1 | none |
| onset | E200K | parent vs. child | 40 | linear regression | 0.68 | 1 | none<sup>32</sup> |
| onset | top three | men vs. women | 446 vs. 492 | Cox | 0.22 | 1 | none |
| duration | top three | men vs. women | 264 vs. 281 | linear regression | 0.02 | 0.44 | yes<sup>5</sup> |
| onset | top three | direct vs. indirect ascertainment | 843 vs. 251 | Cox | 0.78 | 1 | none |
| duration | top three | direct vs. indirect ascertainment | 544 vs. 90 | linear regression | 0.64 | 1 | none |
| onset | top three | year of birth | 697 | linear regression | 3.6E-49 | 7.9E-48 | yes<sup>32</sup> |
| onset | top three | study centers | 973 | two-way ANOVA | 0.012 | 0.26 | none |
| onset | top three | year of onset | 697 | linear regression | 0.00032 | 0.0070 | none |

### Supplementary Life Tables

These tables are made available as .tsv and .xls files in the code and data repository for this manuscript: https://github.com/ericminikel/prnp_onset

### Supplementary Duration Tables

These tables are made available as .tsv and .xls files in the code and data repository for this manuscript: https://github.com/ericminikel/prnp_onset

## Supplementary Figures

**Figure S1.**
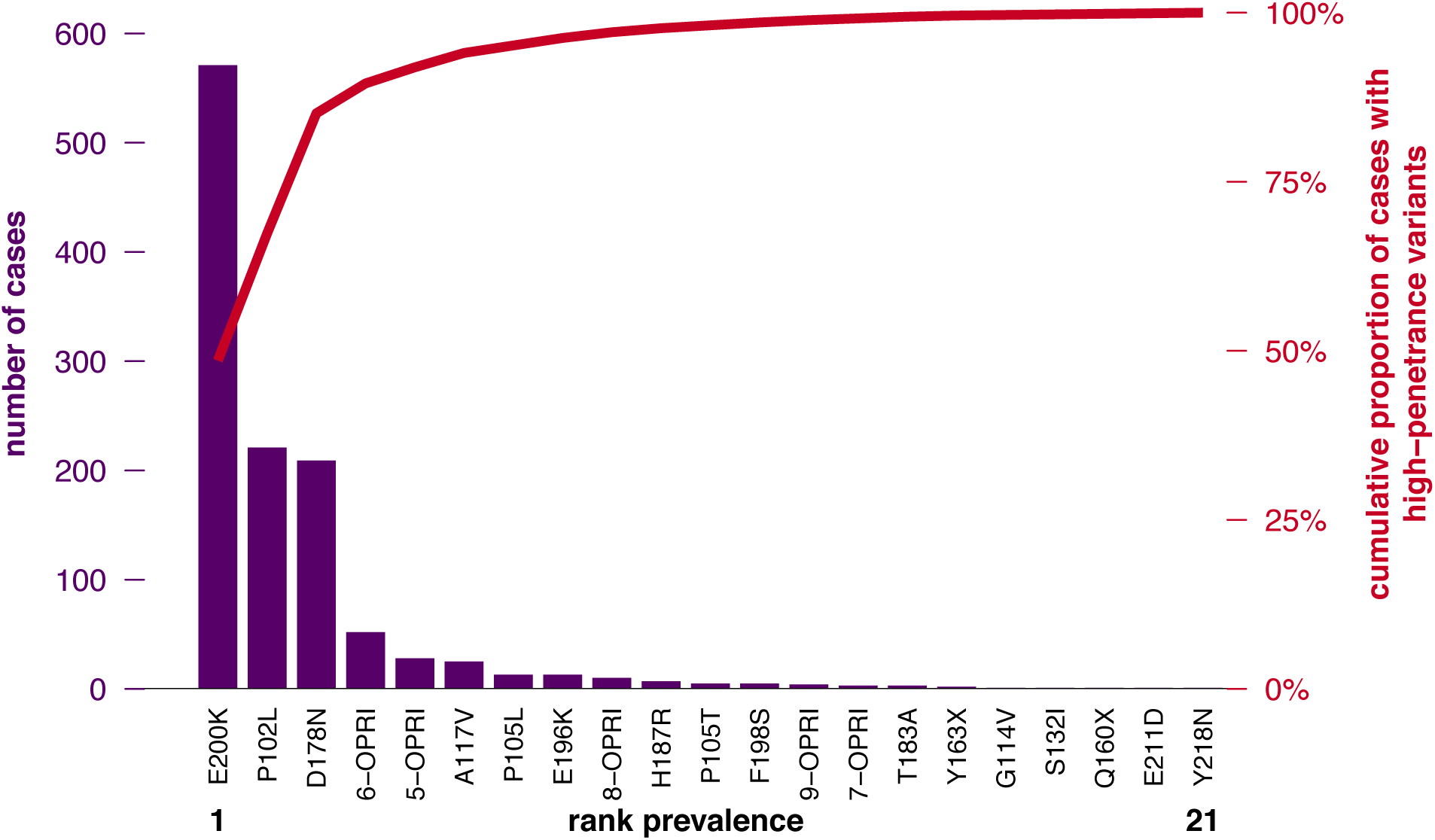
Variant prevalence among prion disease cases with a high penetrance variant. Genetic variants deemed highly penetrant based on the literature review in Table S1 are plotted by the rank (x axis) versus number (left axis) and cumulative proportion (right axis) of high penetrance cases they explain in a recent case series^6^.

**Figure S2.**
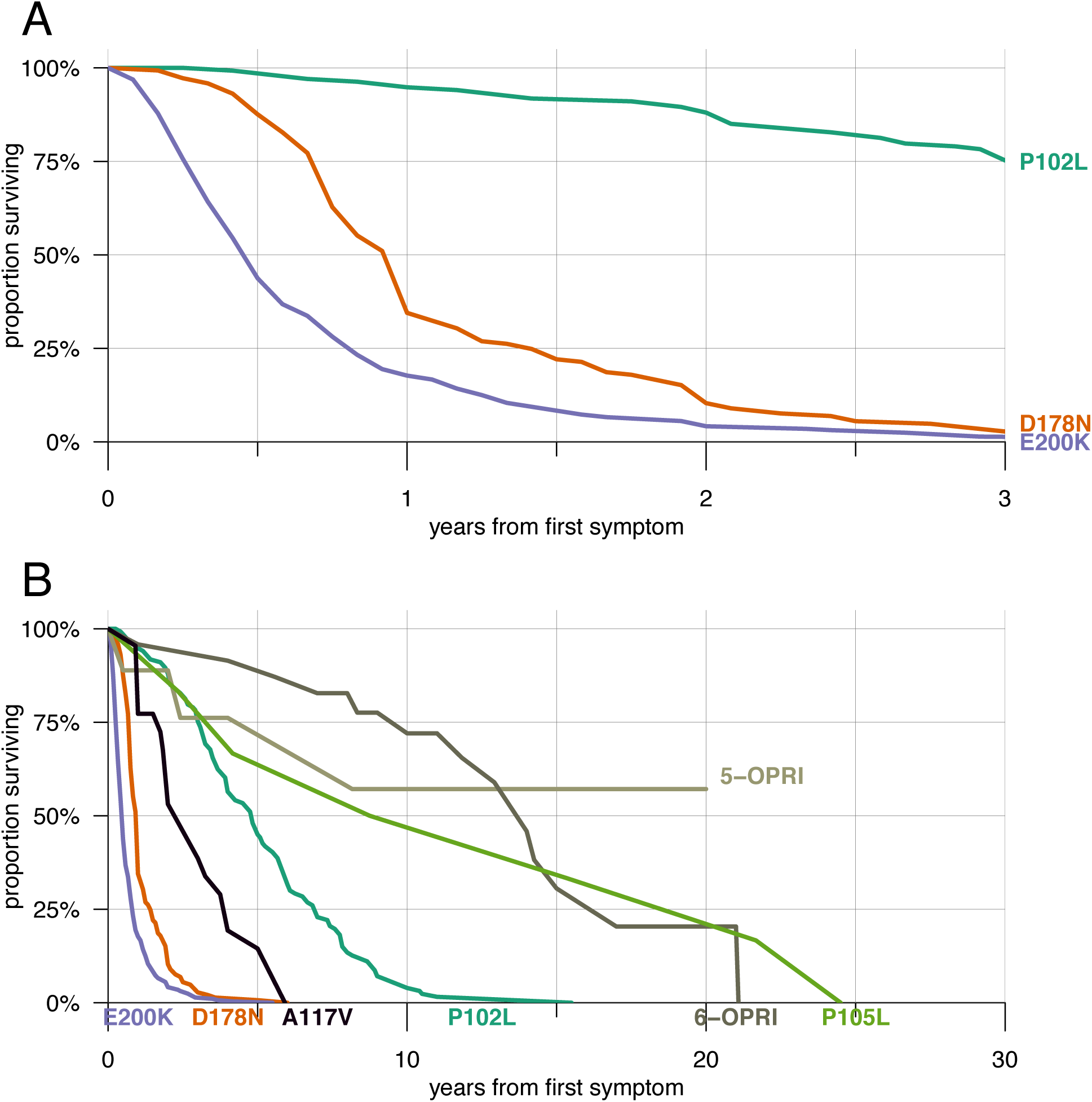
Disease duration by mutation. A) Disease duration (time from first symptom to death) in genetic prion disease. D178N and E200K are classified as rapidly progressive mutations, with >50% of individuals dying within one year of first symptom. B) Zoomed out to 30 years (note y axis) and including supplementary mutations. Disease duration data are provided in the Supplementary Duration Tables.

**Figure S3.**
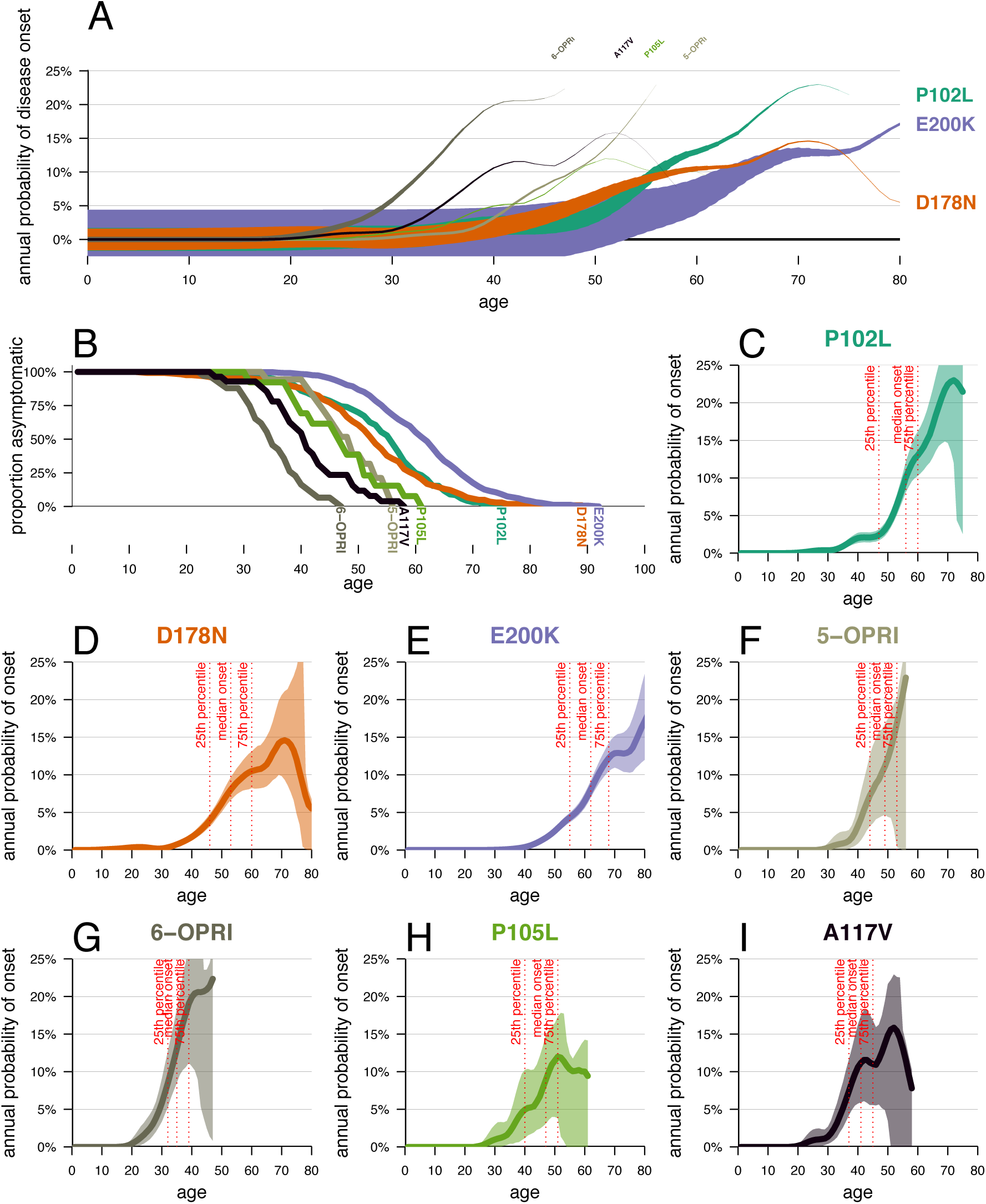
Survival and hazard curves. A) Hazard vs. time with line thickness representing survival, as Figure 1 but including the top 7 mutations. B) Survival curves for the 7 mutations. C-I) Hazard vs. age with 95% confidence intervals displayed in 50% transparency.

**Figure S4.**
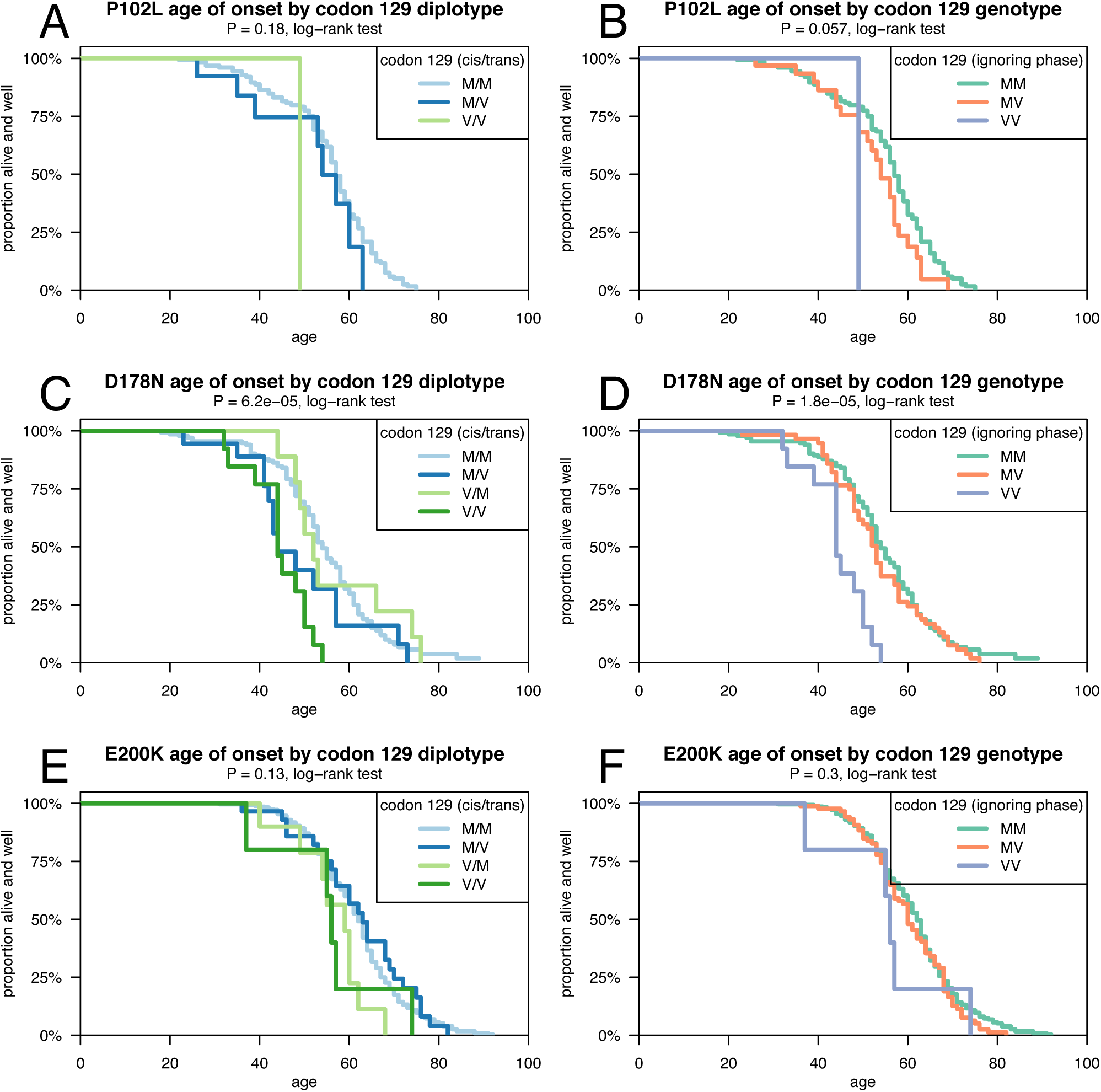
Age of onset and codon 129. Survival curves for age onset or death in P102L (A-B), D178N (C-D), and E200K (E-F) genetic prion disease stratified by codon 129 diplotype (A, C, E) or phaseless genotype (B, D, F).

**Figure S5.**
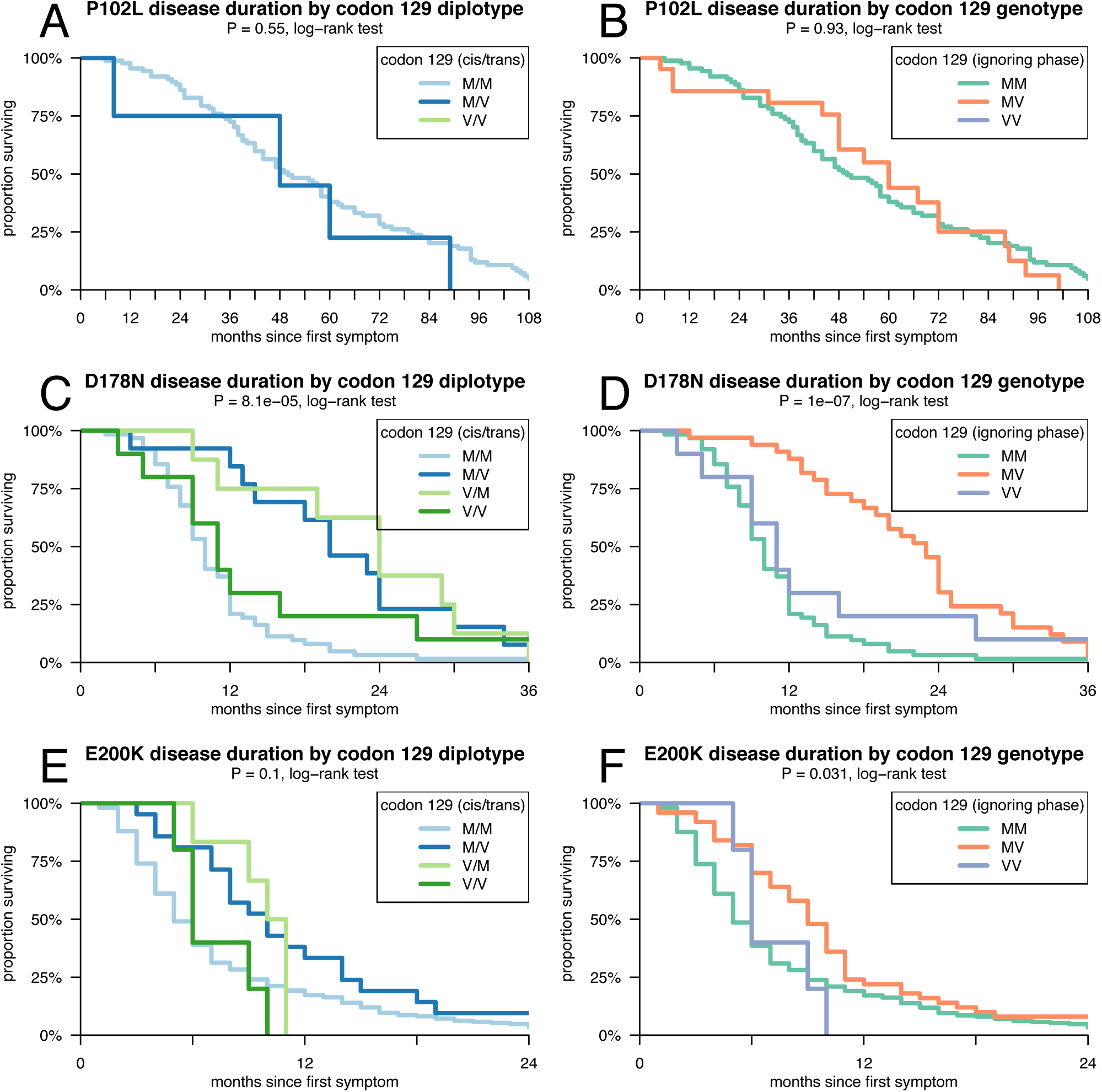
Disease duration and codon 129. Survival curves for disease duration (time from first symptom to death) in P102L (A-B), D178N (C-D), and E200K (E-F) genetic prion disease stratified by codon 129 diplotype (A, C, E) or phaseless genotype (B, D, F).

**Figure S6.**
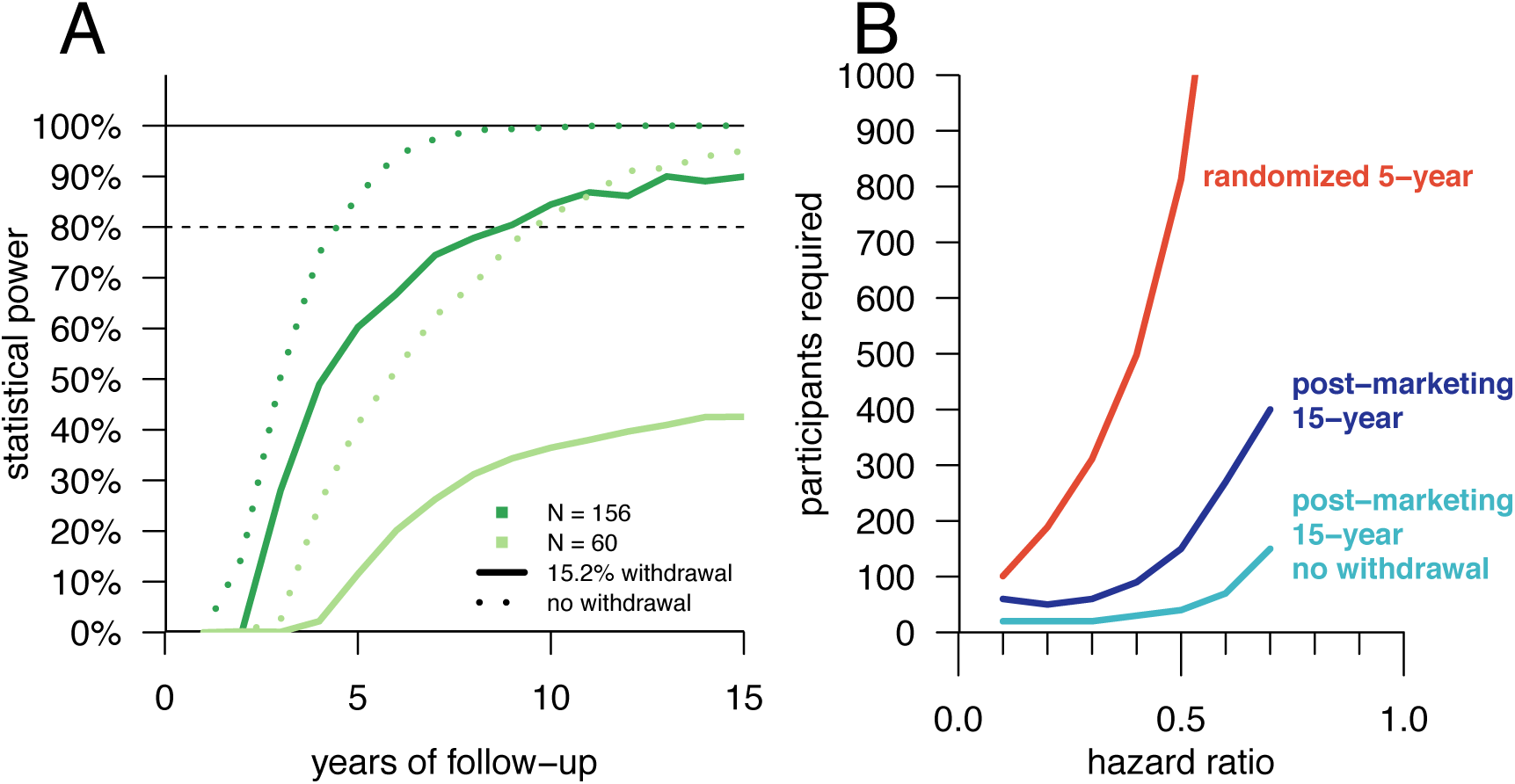
Power increases with long follow-up periods in simulations using historical controls. A) Simulated trial power under the Cox proportional hazards model as a function of the number of individuals randomized and the number of years of follow-up with (solid line) or without (dotted line) modeling withdrawal, assuming a hazard ratio of 0.5 and a run-in period of one year. B) Number of participants required for 80% power at P < 0.05, as a function of hazard ratio (x axis) and trial design (different curves). Numbers for randomized trials (red curve) are taken directly from Table 2, while numbers for post-marketing studies (dark and light blue curves) are obtained by simulation (Supplementary Discussion).

## References

1. New drug, antibiotic, and biological drug product regulations; accelerated approval--FDA. Final rule. Fed Regist. 1992 Dec 11;57(239):58942–58960. PMID: 10123232

2. Naci H, Smalley KR, Kesselheim AS. Characteristics of Preapproval and Postapproval Studies for Drugs Granted Accelerated Approval by the US Food and Drug Administration. JAMA. 2017 Aug 15;318(7):626–636. PMCID: PMC5817559

3. Food and Drug Administration Safety and Innovation Act. Public Law 112-114 Section 506(c)(3). Jul 9, 2012.

4. Prusiner SB. Prions. Proc Natl Acad Sci U S A. 1998 Nov 10;95(23):13363–13383. PMCID: PMC33918

5. Pocchiari M, Puopolo M, Croes EA, Budka H, Gelpi E, Collins S, Lewis V, Sutcliffe T, Guilivi A, Delasnerie-Laupretre N, Brandel J-P, Alperovitch A, Zerr I, Poser S, Kretzschmar HA, Ladogana A, Rietvald I, Mitrova E, Martinez-Martin P, de Pedro-Cuesta J, Glatzel M, Aguzzi A, Cooper S, Mackenzie J, van Duijn CM, Will RG. Predictors of survival in sporadic Creutzfeldt-Jakob disease and other human transmissible spongiform encephalopathies. Brain J Neurol. 2004 Oct;127(Pt 10):2348–2359. PMID: 15361416

6. Minikel EV, Vallabh SM, Lek M, Estrada K, Samocha KE, Sathirapongsasuti JF, McLean CY, Tung JY, Yu LPC, Gambetti P, Blevins J, Zhang S, Cohen Y, Chen W, Yamada M, Hamaguchi T, Sanjo N, Mizusawa H, Nakamura Y, Kitamoto T, Collins SJ, Boyd A, Will RG, Knight R, Ponto C, Zerr I, Kraus TFJ, Eigenbrod S, Giese A, Calero M, de Pedro-Cuesta J, Haïk S, Laplanche J-L, Bouaziz-Amar E, Brandel J-P, Capellari S, Parchi P, Poleggi A, Ladogana A, O’Donnell-Luria AH, Karczewski KJ, Marshall JL, Boehnke M, Laakso M, Mohlke KL, Kähler A, Chambert K, McCarroll S, Sullivan PF, Hultman CM, Purcell SM, Sklar P, van der Lee SJ, Rozemuller A, Jansen C, Hofman A, Kraaij R, van Rooij JGJ, Ikram MA, Uitterlinden AG, van Duijn CM, Exome Aggregation Consortium (ExAC), Daly MJ, MacArthur DG. Quantifying prion disease penetrance using large population control cohorts. Sci Transl Med. 2016 Jan 20;8(322):322ra9. PMID: 26791950

7. Otto M, Cepek L, Ratzka P, Doehlinger S, Boekhoff I, Wiltfang J, Irle E, Pergande G, Ellers-Lenz B, Windl O, Kretzschmar HA, Poser S, Prange H. Efficacy of flupirtine on cognitive function in patients with CJD: A double-blind study. Neurology. 2004 Mar 9;62(5):714–718. PMID: 15007119

8. Bone I, Belton L, Walker AS, Darbyshire J. Intraventricular pentosan polysulphate in human prion diseases: an observational study in the UK. Eur J Neurol. 2008 May;15(5):458–464. PMID: 18355301

9. Tsuboi Y, Doh-Ura K, Yamada T. Continuous intraventricular infusion of pentosan polysulfate: clinical trial against prion diseases. Neuropathol Off J Jpn Soc Neuropathol. 2009 Oct;29(5):632–636. PMID: 19788637

10. Haïk S, Brandel JP, Salomon D, Sazdovitch V, Delasnerie-Lauprêtre N, Laplanche JL, Faucheux BA, Soubrié C, Boher E, Belorgey C, Hauw JJ, Alpérovitch A. Compassionate use of quinacrine in Creutzfeldt-Jakob disease fails to show significant effects. Neurology. 2004 Dec 28;63(12):2413–2415. PMID: 15623716

11. Collinge J, Gorham M, Hudson F, Kennedy A, Keogh G, Pal S, Rossor M, Rudge P, Siddique D, Spyer M, Thomas D, Walker S, Webb T, Wroe S, Darbyshire J. Safety and efficacy of quinacrine in human prion disease (PRION-1 study): a patient-preference trial. Lancet Neurol. 2009 Apr;8(4):334–344. PMCID: PMC2660392

12. Geschwind MD, Kuo AL, Wong KS, Haman A, Devereux G, Raudabaugh BJ, Johnson DY, Torres-Chae CC, Finley R, Garcia P, Thai JN, Cheng HQ, Neuhaus JM, Forner SA, Duncan JL, Possin KL, Dearmond SJ, Prusiner SB, Miller BL. Quinacrine treatment trial for sporadic Creutzfeldt-Jakob disease. Neurology. 2013 Dec 3;81(23):2015–2023. PMCID: PMC4211922

13. Haïk S, Marcon G, Mallet A, Tettamanti M, Welaratne A, Giaccone G, Azimi S, Pietrini V, Fabreguettes J-R, Imperiale D, Cesaro P, Buffa C, Aucan C, Lucca U, Peckeu L, Suardi S, Tranchant C, Zerr I, Houillier C, Redaelli V, Vespignani H, Campanella A, Sellal F, Krasnianski A, Seilhean D, Heinemann U, Sedel F, Canovi M, Gobbi M, Di Fede G, Laplanche J-L, Pocchiari M, Salmona M, Forloni G, Brandel J-P, Tagliavini F. Doxycycline in Creutzfeldt-Jakob disease: a phase 2, randomised, double-blind, placebo-controlled trial. Lancet Neurol. 2014 Feb;13(2):150–158. PMID: 24411709

14. Varges D, Manthey H, Heinemann U, Ponto C, Schmitz M, Schulz-Schaeffer WJ, Krasnianski A, Breithaupt M, Fincke F, Kramer K, Friede T, Zerr I. Doxycycline in early CJD: a double-blinded randomised phase II and observational study. J Neurol Neurosurg Psychiatry. 2017 Feb;88(2):119–125. PMCID: PMC5284486

15. Bechtel K, Geschwind MD. Ethics in prion disease. Prog Neurobiol. 2013 Nov;110:29–44. PMCID: PMC3818451

16. Doh-ura K, Ishikawa K, Murakami-Kubo I, Sasaki K, Mohri S, Race R, Iwaki T. Treatment of transmissible spongiform encephalopathy by intraventricular drug infusion in animal models. J Virol. 2004 May;78(10):4999–5006. PMCID: PMC400350

17. Kawasaki Y, Kawagoe K, Chen C, Teruya K, Sakasegawa Y, Doh-ura K. Orally administered amyloidophilic compound is effective in prolonging the incubation periods of animals cerebrally infected with prion diseases in a prion strain-dependent manner. J Virol. 2007 Dec;81(23):12889–12898. PMCID: PMC2169081

18. Wagner J, Ryazanov S, Leonov A, Levin J, Shi S, Schmidt F, Prix C, Pan-Montojo F, Bertsch U, Mitteregger-Kretzschmar G, Geissen M, Eiden M, Leidel F, Hirschberger T, Deeg AA, Krauth JJ, Zinth W, Tavan P, Pilger J, Zweckstetter M, Frank T, Bähr M, Weishaupt JH, Uhr M, Urlaub H, Teichmann U, Samwer M, Bötzel K, Groschup M, Kretzschmar H, Griesinger C, Giese A. Anle138b: a novel oligomer modulator for disease-modifying therapy of neurodegenerative diseases such as prion and Parkinson’s disease. Acta Neuropathol (Berl). 2013 Jun;125(6):795–813. PMCID: PMC3661926

19. Giles K, Berry DB, Condello C, Hawley RC, Gallardo-Godoy A, Bryant C, Oehler A, Elepano M, Bhardwaj S, Patel S, Silber BM, Guan S, DeArmond SJ, Renslo AR, Prusiner SB. Different 2- Aminothiazole Therapeutics Produce Distinct Patterns of Scrapie Prion Neuropathology in Mouse Brains. J Pharmacol Exp Ther. 2015 Oct;355(1):2–12. PMID: 26224882

20. Tariot PN, Lopera F, Langbaum JB, Thomas RG, Hendrix S, Schneider LS, Rios-Romenets S, Giraldo M, Acosta N, Tobon C, Ramos C, Espinosa A, Cho W, Ward M, Clayton D, Friesenhahn M, Mackey H, Honigberg L, Sanabria Bohorquez S, Chen K, Walsh T, Langlois C, Reiman EM, Alzheimer’s Prevention Initiative. The Alzheimer’s Prevention Initiative Autosomal-Dominant Alzheimer’s Disease Trial: A study of crenezumab versus placebo in preclinical PSEN1 E280A mutation carriers to evaluate efficacy and safety in the treatment of autosomal-dominant Alzheimer’s disease, including a placebo-treated noncarrier cohort. Alzheimers Dement N Y N. 2018;4:150–160. PMCID: PMC6021543

21. Kovács GG, Puopolo M, Ladogana A, Pocchiari M, Budka H, van Duijn C, Collins SJ, Boyd A, Giulivi A, Coulthart M, Delasnerie-Laupretre N, Brandel JP, Zerr I, Kretzschmar HA, de Pedro-Cuesta J, Calero-Lara M, Glatzel M, Aguzzi A, Bishop M, Knight R, Belay G, Will R, Mitrova E, EUROCJD. Genetic prion disease: the EUROCJD experience. Hum Genet. 2005 Nov;118(2):166–174. PMID: 16187142

22. Nozaki I, Hamaguchi T, Sanjo N, Noguchi-Shinohara M, Sakai K, Nakamura Y, Sato T, Kitamoto T, Mizusawa H, Moriwaka F, Shiga Y, Kuroiwa Y, Nishizawa M, Kuzuhara S, Inuzuka T, Takeda M, Kuroda S, Abe K, Murai H, Murayama S, Tateishi J, Takumi I, Shirabe S, Harada M, Sadakane A, Yamada M. Prospective 10-year surveillance of human prion diseases in Japan. Brain J Neurol. 2010 Oct;133(10):3043–3057. PMID: 20855418

23. Owen J, Beck J, Campbell T, Adamson G, Gorham M, Thompson A, Smithson S, Rosser E, Rudge P, Collinge J, Mead S. Predictive testing for inherited prion disease: report of 22 years experience. Eur J Hum Genet EJHG. 2014 Apr 9; PMID: 24713662

24. Cooper DN, Youssoufian H. The CpG dinucleotide and human genetic disease. Hum Genet. 1988 Feb;78(2):151–155. PMID: 3338800

25. Samocha KE, Robinson EB, Sanders SJ, Stevens C, Sabo A, McGrath LM, Kosmicki JA, Rehnström K, Mallick S, Kirby A, Wall DP, MacArthur DG, Gabriel SB, DePristo M, Purcell SM, Palotie A, Boerwinkle E, Buxbaum JD, Cook EH, Gibbs RA, Schellenberg GD, Sutcliffe JS, Devlin B, Roeder K, Neale BM, Daly MJ. A framework for the interpretation of de novo mutation in human disease. Nat Genet. 2014 Sep;46(9):944–950. PMCID: PMC4222185

26. Lee HS, Sambuughin N, Cervenakova L, Chapman J, Pocchiari M, Litvak S, Qi HY, Budka H, del Ser T, Furukawa H, Brown P, Gajdusek DC, Long JC, Korczyn AD, Goldfarb LG. Ancestral origins and worldwide distribution of the PRNP 200K mutation causing familial Creutzfeldt-Jakob disease. Am J Hum Genet. 1999 Apr;64(4):1063–1070. PMCID: PMC1377830

27. Dagvadorj A, Petersen RB, Lee HS, Cervenakova L, Shatunov A, Budka H, Brown P, Gambetti P, Goldfarb LG. Spontaneous mutations in the prion protein gene causing transmissible spongiform encephalopathy. Ann Neurol. 2002 Sep;52(3):355–359. PMID: 12205650

28. Kong Q, Surewicz WK, Petersen RB, Chen SG, Gambetti P, Parchi P, Capellari S, Goldfarb L, Montagna P, Lugaresi E, Piccardo P, Ghetti B. Inherited Prion Diseases. Prion Biol Dis [Internet]. 2nd ed. Cold Spring Harbor Laboratory Press; 2004. Available from: https://cshmonographs.org/index.php/monographs/article/viewArticle/4035

29. Klug GMJA, Wand H, Simpson M, Boyd A, Law M, Masters CL, Matěj R, Howley R, Farrell M, Breithaupt M, Zerr I, van Duijn C, Ibrahim-Verbaas C, Mackenzie J, Will RG, Brandel J-P, Alperovitch A, Budka H, Kovacs GG, Jansen GH, Coulthard M, Collins SJ. Intensity of human prion disease surveillance predicts observed disease incidence. J Neurol Neurosurg Psychiatry. 2013 Dec;84(12):1372–1377. PMID: 23965290

30. Schoenfeld DA. Sample-size formula for the proportional-hazards regression model. Biometrics. 1983 Jun;39(2):499–503. PMID: 6354290

31. Woodcock J. Reforming Clinical Trials in Drug Development: Impact of Targeted Therapies [Internet]. 2016 Nov 16. Available from: https://www.fda.gov/downloads/AboutFDA/CentersOffices/OfficeofMedicalProductsandTobacco/C DER/UCM530409.pdf

32. Minikel EV, Zerr I, Collins SJ, Ponto C, Boyd A, Klug G, Karch A, Kenny J, Collinge J, Takada LT, Forner S, Fong JC, Mead S, Geschwind MD. Ascertainment bias causes false signal of anticipation in genetic prion disease. Am J Hum Genet. 2014 Oct 2;95(4):371–382. PMCID: PMC4185115

33. Capellari S, Strammiello R, Saverioni D, Kretzschmar H, Parchi P. Genetic Creutzfeldt-Jakob disease and fatal familial insomnia: insights into phenotypic variability and disease pathogenesis. Acta Neuropathol (Berl). 2011 Jan;121(1):21–37. PMID: 20978903

34. Mead S. Prion disease genetics. Eur J Hum Genet EJHG. 2006 Mar;14(3):273–281. PMID: 16391566

35. Mead S, Poulter M, Beck J, Webb TEF, Campbell TA, Linehan JM, Desbruslais M, Joiner S, Wadsworth JDF, King A, Lantos P, Collinge J. Inherited prion disease with six octapeptide repeat insertional mutation--molecular analysis of phenotypic heterogeneity. Brain J Neurol. 2006 Sep;129(Pt 9):2297–2317. PMID: 16923955

36. Webb TEF, Poulter M, Beck J, Uphill J, Adamson G, Campbell T, Linehan J, Powell C, Brandner S, Pal S, Siddique D, Wadsworth JD, Joiner S, Alner K, Petersen C, Hampson S, Rhymes C, Treacy C, Storey E, Geschwind MD, Nemeth AH, Wroe S, Collinge J, Mead S. Phenotypic heterogeneity and genetic modification of P102L inherited prion disease in an international series. Brain J Neurol. 2008 Oct;131(Pt 10):2632–2646. PMCID: PMC2570713

37. Lerman C, Narod S, Schulman K, Hughes C, Gomez-Caminero A, Bonney G, Gold K, Trock B, Main D, Lynch J, Fulmore C, Snyder C, Lemon SJ, Conway T, Tonin P, Lenoir G, Lynch H. BRCA1 testing in families with hereditary breast-ovarian cancer. A prospective study of patient decision making and outcomes. JAMA. 1996 Jun 26;275(24):1885–1892. PMID: 8648868

38. Green RC, Berg JS, Grody WW, Kalia SS, Korf BR, Martin CL, McGuire AL, Nussbaum RL, O’Daniel JM, Ormond KE, Rehm HL, Watson MS, Williams MS, Biesecker LG, American College of Medical Genetics and Genomics. ACMG recommendations for reporting of incidental findings in clinical exome and genome sequencing. Genet Med Off J Am Coll Med Genet. 2013 Jul;15(7):565–574. PMCID: PMC3727274

39. U.S. Food and Drug Administration. Rare Diseases: Common Issues in Drug Development. Draft Guidance for Industry. [Internet]. 2015 [cited 2018 Aug 22]. Available from: https://www.fda.gov/downloads/Drugs/GuidanceComplianceRegulatoryInformation/Guidances/uc m458485.pdf

40. Hamasaki S, Shirabe S, Tsuda R, Yoshimura T, Nakamura T, Eguchi K. Discordant Gerstmann-Sträussler-Scheinker disease in monozygotic twins. Lancet Lond Engl. 1998 Oct 24;352(9137):1358–1359. PMID: 9802281

41. Mead S, Uphill J, Beck J, Poulter M, Campbell T, Lowe J, Adamson G, Hummerich H, Klopp N, Rückert I-M, Wichmann H-E, Azazi D, Plagnol V, Pako WH, Whitfield J, Alpers MP, Whittaker J, Balding DJ, Zerr I, Kretzschmar H, Collinge J. Genome-wide association study in multiple human prion diseases suggests genetic risk factors additional to PRNP. Hum Mol Genet. 2012 Apr 15;21(8):1897–1906. PMCID: PMC3313791

42. Lopera F, Ardilla A, Martínez A, Madrigal L, Arango-Viana JC, Lemere CA, Arango-Lasprilla JC, Hincapíe L, Arcos-Burgos M, Ossa JE, Behrens IM, Norton J, Lendon C, Goate AM, Ruiz-Linares A, Rosselli M, Kosik KS. Clinical features of early-onset Alzheimer disease in a large kindred with an E280A presenilin-1 mutation. JAMA. 1997 Mar 12;277(10):793–799. PMID: 9052708

43. Ryman DC, Acosta-Baena N, Aisen PS, Bird T, Danek A, Fox NC, Goate A, Frommelt P, Ghetti B, Langbaum JBS, Lopera F, Martins R, Masters CL, Mayeux RP, McDade E, Moreno S, Reiman EM, Ringman JM, Salloway S, Schofield PR, Sperling R, Tariot PN, Xiong C, Morris JC, Bateman RJ, and the Dominantly Inherited Alzheimer Network. Symptom onset in autosomal dominant Alzheimer disease: A systematic review and meta-analysis. Neurology. 2014 Jun 13; PMID: 24928124

44. Mullard A. Sting of Alzheimer’s failures offset by upcoming prevention trials. Nat Rev Drug Discov. 2012 Sep;11(9):657–660. PMID: 22935790

45. Reiman EM, Langbaum JBS, Fleisher AS, Caselli RJ, Chen K, Ayutyanont N, Quiroz YT, Kosik KS, Lopera F, Tariot PN. Alzheimer’s Prevention Initiative: a plan to accelerate the evaluation of presymptomatic treatments. J Alzheimers Dis JAD. 2011;26 Suppl 3:321–329. PMCID: PMC3343739

46. The Colombian Alzheimer’s Prevention Initiative (API) Registry [Internet]. [cited 2017 Mar 22]. Available from: http://www.sciencedirect.com/science/article/pii/S155252601632965X

47. Garber K. Genentech’s Alzheimer’s antibody trial to study disease prevention. Nat Biotechnol. 2012 Aug;30(8):731–732. PMID: 22871696

48. McDade E, Bateman RJ. Stop Alzheimer’s before it starts. Nature. 2017 12;547(7662):153–155. PMID: 28703214

49. Vallabh SM, Nobuhara CK, Llorens F, Zerr I, Parchi P, Capellari S, Kuhn E, Klickstein J, Safar J, Nery F, Swoboda K, Schreiber SL, Geschwind MD, Zetterberg H, Arnold SE, Minikel EV. Prion protein quantification in cerebrospinal fluid as a tool for prion disease drug development. bioRxiv [Internet]. 2018 Jan 1; Available from: http://biorxiv.org/content/early/2018/04/04/295063.abstract

50. Vallabh S, Minikel EV, Schreiber SL, Lander ES. A path to prevention of genetic prion disease. In preparation.

51. Brandenburg NA, Bwire R, Freeman J, Houn F, Sheehan P, Zeldis JB. Effectiveness of Risk Evaluation and Mitigation Strategies (REMS) for Lenalidomide and Thalidomide: Patient Comprehension and Knowledge Retention. Drug Saf. 2017 Apr;40(4):333–341. PMID: 28074423

52. Reynolds IS, Rising JP, Coukell AJ, Paulson KH, Redberg RF. Assessing the safety and effectiveness of devices after US Food and Drug Administration approval: FDA-mandated postapproval studies. JAMA Intern Med. 2014 Nov;174(11):1773–1779. PMID: 25265209

53. Rathi VK, Krumholz HM, Masoudi FA, Ross JS. Characteristics of Clinical Studies Conducted Over the Total Product Life Cycle of High-Risk Therapeutic Medical Devices Receiving FDA Premarket Approval in 2010 and 2011. JAMA. 2015 Aug 11;314(6):604–612. PMID: 26262798

54. Langbehn DR, Hayden MR, Paulsen JS, PREDICT-HD Investigators of the Huntington Study Group. CAG-repeat length and the age of onset in Huntington disease (HD): a review and validation study of statistical approaches. Am J Med Genet Part B Neuropsychiatr Genet Off Publ Int Soc Psychiatr Genet. 2010 Mar 5;153B(2):397–408. PMCID: PMC3048807

55. Langbehn DR, Brinkman RR, Falush D, Paulsen JS, Hayden MR, International Huntington’s Disease Collaborative Group. A new model for prediction of the age of onset and penetrance for Huntington’s disease based on CAG length. Clin Genet. 2004 Apr;65(4):267–277. PMID: 15025718

56. Collins S, Boyd A, Lee JS, Lewis V, Fletcher A, McLean CA, Law M, Kaldor J, Smith MJ, Masters CL. Creutzfeldt-Jakob disease in Australia 1970-1999. Neurology. 2002 Nov 12;59(9):1365–1371. PMID: 12427885

57. Windl O, Giese A, Schulz-Schaeffer W, Zerr I, Skworc K, Arendt S, Oberdieck C, Bodemer M, Poser S, Kretzschmar HA. Molecular genetics of human prion diseases in Germany. Hum Genet. 1999 Sep;105(3):244–252. PMID: 10987652

58. Beck JA, Poulter M, Campbell TA, Adamson G, Uphill JB, Guerreiro R, Jackson GS, Stevens JC, Manji H, Collinge J, Mead S. PRNP allelic series from 19 years of prion protein gene sequencing at the MRC Prion Unit. Hum Mutat. 2010 Jul;31(7):E1551–1563. PMID: 20583301

59. Takada LT, Kim M-O, Cleveland RW, Wong K, Forner SA, Gala II, Fong JC, Geschwind MD. Genetic prion disease: Experience of a rapidly progressive dementia center in the United States and a review of the literature. Am J Med Genet Part B Neuropsychiatr Genet Off Publ Int Soc Psychiatr Genet. 2017 Jan;174(1):36–69. PMID: 27943639

60. Forloni G, Tettamanti M, Lucca U, Albanese Y, Quaglio E, Chiesa R, Erbetta A, Villani F, Redaelli V, Tagliavini F, Artuso V, Roiter I. Preventive study in subjects at risk of fatal familial insomnia: Innovative approach to rare diseases. Prion. 2015;9(2):75–79. PMCID: PMC4601344

61. Parchi P, Giese A, Capellari S, Brown P, Schulz-Schaeffer W, Windl O, Zerr I, Budka H, Kopp N, Piccardo P, Poser S, Rojiani A, Streichemberger N, Julien J, Vital C, Ghetti B, Gambetti P, Kretzschmar H. Classification of sporadic Creutzfeldt-Jakob disease based on molecular and phenotypic analysis of 300 subjects. Ann Neurol. 1999 Aug;46(2):224–233. PMID: 10443888

62. Michael Kohn, Josh Senyak, Mike Jarrett. Sample Size Calculators, UCSF Clinical & Translational Sciences Institute [Internet]. 2017 [cited 2017 Feb 9]. Available from: http://www.sample-size.net/sample-size-survival-analysis/

63. Lek M, Karczewski KJ, Minikel EV, Samocha KE, Banks E, Fennell T, O’Donnell-Luria AH, Ware JS, Hill AJ, Cummings BB, Tukiainen T, Birnbaum DP, Kosmicki JA, Duncan LE, Estrada K, Zhao F, Zou J, Pierce-Hoffman E, Berghout J, Cooper DN, Deflaux N, DePristo M, Do R, Flannick J, Fromer M, Gauthier L, Goldstein J, Gupta N, Howrigan D, Kiezun A, Kurki MI, Moonshine AL, Natarajan P, Orozco L, Peloso GM, Poplin R, Rivas MA, Ruano-Rubio V, Rose SA, Ruderfer DM, Shakir K, Stenson PD, Stevens C, Thomas BP, Tiao G, Tusie-Luna MT, Weisburd B, Won H-H, Yu D, Altshuler DM, Ardissino D, Boehnke M, Danesh J, Donnelly S, Elosua R, Florez JC, Gabriel SB, Getz G, Glatt SJ, Hultman CM, Kathiresan S, Laakso M, McCarroll S, McCarthy MI, McGovern D, McPherson R, Neale BM, Palotie A, Purcell SM, Saleheen D, Scharf JM, Sklar P, Sullivan PF, Tuomilehto J, Tsuang MT, Watkins HC, Wilson JG, Daly MJ, MacArthur DG, Exome Aggregation Consortium. Analysis of protein-coding genetic variation in 60,706 humans. Nature. 2016 Aug 18;536(7616):285–291. PMCID: PMC5018207

64. Mitrová E, Belay G. Creutzfeldt-Jakob disease with E200K mutation in Slovakia: characterization and development. Acta Virol. 2002;46(1):31–39. PMID: 12197632

65. Cohen OS, Chapman J, Korczyn AD, Nitsan Z, Appel S, Hoffmann C, Rosenmann H, Kahana E, Lee H. Familial Creutzfeldt-Jakob disease with the E200K mutation: longitudinal neuroimaging from asymptomatic to symptomatic CJD. J Neurol. 2015 Mar;262(3):604–613. PMID: 25522698

66. Murray K. Creutzfeldt-Jacob disease mimics, or how to sort out the subacute encephalopathy patient. Pract Neurol. 2011 Feb;11(1):19–28. PMID: 21239650

67. Morrison PJ, Harding-Lester S, Bradley A. Uptake of Huntington disease predictive testing in a complete population. Clin Genet. 2011 Sep;80(3):281–286. PMID: 20880124

68. U.S. Food and Drug Administration. Small Business Assistance: Frequently Asked Questions on the Patent Term Restoration Program [Internet]. 2017 [cited 2018 Aug 22]. Available from: https://www.fda.gov/Drugs/DevelopmentApprovalProcess/SmallBusinessAssistance/ucm069959. htm

69. U.S. Food and Drug Administration. Reference Product Exclusivity for Biological Products Filed Under Section 351(a) of the PHS Act. Draft Guidance for Industry. [Internet]. 2014 [cited 2018 Aug 24]. Available from: https://www.fda.gov/downloads/drugs/guidancecomplianceregulatoryinformation/guidances/ucm4 07844.pdf

70. U.S. Food and Drug Administration. Frequently Asked Questions on Patents and Exclusivity [Internet]. 2018 [cited 2018 Aug 22]. Available from: https://www.fda.gov/drugs/developmentapprovalprocess/ucm079031.htm

71. Wang B, Liu J, Kesselheim AS. Variations in time of market exclusivity among top-selling prescription drugs in the United States. JAMA Intern Med. 2015 Apr;175(4):635–637. PMID: 25664700

72. Grabowski H, Long G, Mortimer R. Recent trends in brand-name and generic drug competition. J Med Econ. 2014 Mar;17(3):207–214. PMID: 24320785

73. Downing NS, Aminawung JA, Shah ND, Krumholz HM, Ross JS. Clinical trial evidence supporting FDA approval of novel therapeutic agents, 2005-2012. JAMA. 2014 Jan 22;311(4):368–377. PMCID: PMC4144867

74. Klein JP, Moeschberger ML. Survival Analysis - Techniques for Censored and Truncated Data [Internet]. 2003 [cited 2017 Oct 3]. Available from: http://www.springer.com/us/book/9780387953991

75. Bernardi L, Cupidi C, Frangipane F, Anfossi M, Gallo M, Conidi ME, Vasso F, Colao R, Puccio G, Curcio SAM, Mirabelli M, Clodomiro A, Di Lorenzo R, Smirne N, Maletta R, Bruni AC. Novel N-terminal domain mutation in prion protein detected in 2 patients diagnosed with frontotemporal lobar degeneration syndrome. Neurobiol Aging. 2014 Nov;35(11):2657.e7–11. PMID: 25022973

76. Beck JA, Mead S, Campbell TA, Dickinson A, Wientjens DP, Croes EA, Van Duijn CM, Collinge J. Two-octapeptide repeat deletion of prion protein associated with rapidly progressive dementia. Neurology. 2001 Jul 24;57(2):354–356. PMID: 11468331

77. Capellari S, Parchi P, Wolff BD, Campbell J, Atkinson R, Posey DM, Petersen RB, Gambetti P. Creutzfeldt-Jakob disease associated with a deletion of two repeats in the prion protein gene. Neurology. 2002 Nov 26;59(10):1628–1630. PMID: 12451210

78. Laplanche JL, Delasnerie-Lauprêtre N, Brandel JP, Dussaucy M, Chatelain J, Launay JM. Two novel insertions in the prion protein gene in patients with late-onset dementia. Hum Mol Genet. 1995 Jun;4(6):1109–1111. PMID: 7655470

79. Pietrini V, Puoti G, Limido L, Rossi G, Di Fede G, Giaccone G, Mangieri M, Tedeschi F, Bondavalli A, Mancia D, Bugiani O, Tagliavini F. Creutzfeldt-Jakob disease with a novel extra-repeat insertional mutation in the PRNP gene. Neurology. 2003 Nov 11;61(9):1288–1291. PMID: 14610142

80. Hill AF, Joiner S, Beck JA, Campbell TA, Dickinson A, Poulter M, Wadsworth JDF, Collinge J. Distinct glycoform ratios of protease resistant prion protein associated with PRNP point mutations. Brain J Neurol. 2006 Mar;129(Pt 3):676–685. PMID: 16415305

81. Nishida Y, Sodeyama N, Toru Y, Toru S, Kitamoto T, Mizusawa H. Creutzfeldt-Jakob disease with a novel insertion and codon 219 Lys/Lys polymorphism in PRNP. Neurology. 2004 Nov 23;63(10):1978–1979. PMID: 15557533

82. Kaski DN, Pennington C, Beck J, Poulter M, Uphill J, Bishop MT, Linehan JM, O’Malley C, Wadsworth JDF, Joiner S, Knight RSG, Ironside JW, Brandner S, Collinge J, Mead S. Inherited prion disease with 4-octapeptide repeat insertion: disease requires the interaction of multiple genetic risk factors. Brain J Neurol. 2011 Jun;134(Pt 6):1829–1838. PMID: 21616973

83. Mead S, Webb TEF, Campbell TA, Beck J, Linehan JM, Rutherfoord S, Joiner S, Wadsworth JDF, Heckmann J, Wroe S, Doey L, King A, Collinge J. Inherited prion disease with 5-OPRI: phenotype modification by repeat length and codon 129. Neurology. 2007 Aug 21;69(8):730–738. PMID: 17709704

84. Goldfarb LG, Brown P, McCombie WR, Goldgaber D, Swergold GD, Wills PR, Cervenakova L, Baron H, Gibbs CJ, Gajdusek DC. Transmissible familial Creutzfeldt-Jakob disease associated with five, seven, and eight extra octapeptide coding repeats in the PRNP gene. Proc Natl Acad Sci U S A. 1991 Dec 1;88(23):10926–10930. PMCID: PMC53045

85. Laplanche JL, Hachimi KH, Durieux I, Thuillet P, Defebvre L, Delasnerie-Lauprêtre N, Peoc’h K, Foncin JF, Destée A. Prominent psychiatric features and early onset in an inherited prion disease with a new insertional mutation in the prion protein gene. Brain J Neurol. 1999 Dec;122 (Pt 12):2375–2386. PMID: 10581230

86. Krasemann S, Zerr I, Weber T, Poser S, Kretzschmar H, Hunsmann G, Bodemer W. Prion disease associated with a novel nine octapeptide repeat insertion in the PRNP gene. Brain Res Mol Brain Res. 1995 Dec 1;34(1):173–176. PMID: 8750875

87. Sánchez-Valle R, Aróstegui JI, Yagüe J, Rami L, Lladó A, Molinuevo JL. First demonstrated de novo insertion in the prion protein gene in a young patient with dementia. J Neurol Neurosurg Psychiatry. 2008 Jul;79(7):845–846. PMID: 18559465

88. Kumar N, Boeve BF, Boot BP, Orr CF, Duffy J, Woodruff BK, Nair AK, Ellison J, Kuntz K, Kantarci K, Jack CR, Westmoreland BF, Fields JA, Baker M, Rademakers R, Parisi JE, Dickson DW. Clinical characterization of a kindred with a novel 12-octapeptide repeat insertion in the prion protein gene. Arch Neurol. 2011 Sep;68(9):1165–1170. PMCID: PMC3326586

89. Jones M, Odunsi S, du Plessis D, Vincent A, Bishop M, Head MW, Ironside JW, Gow D. Gerstmann-Straüssler-Scheinker disease: novel PRNP mutation and VGKC-complex antibodies. Neurology. 2014 Jun 10;82(23):2107–2111. PMCID: PMC4118501

90. Zheng L, Longfei J, Jing Y, Xinqing Z, Haiqing S, Haiyan L, Fen W, Xiumin D, Jianping J. PRNP mutations in a series of apparently sporadic neurodegenerative dementias in China. Am J Med Genet Part B Neuropsychiatr Genet Off Publ Int Soc Psychiatr Genet. 2008 Sep 5;147B(6):938–944. PMID: 18425766

91. Yamada M, Itoh Y, Inaba A, Wada Y, Takashima M, Satoh S, Kamata T, Okeda R, Kayano T, Suematsu N, Kitamoto T, Otomo E, Matsushita M, Mizusawa H. An inherited prion disease with a PrP P105L mutation: clinicopathologic and PrP heterogeneity. Neurology. 1999 Jul 13;53(1):181–188. PMID: 10408557

92. Tunnell E, Wollman R, Mallik S, Cortes CJ, Dearmond SJ, Mastrianni JA. A novel PRNP-P105S mutation associated with atypical prion disease and a rare PrPSc conformation. Neurology. 2008 Oct 28;71(18):1431–1438. PMCID: PMC2676963

93. Rogaeva E, Zadikoff C, Ponesse J, Schmitt-Ulms G, Kawarai T, Sato C, Salehi-Rad S, St George-Hyslop P, Lang AE. Childhood onset in familial prion disease with a novel mutation in the PRNP gene. Arch Neurol. 2006 Jul;63(7):1016–1021. PMID: 16831973

94. Rodriguez M-M, Peoc’h K, Haïk S, Bouchet C, Vernengo L, Mañana G, Salamano R, Carrasco L, Lenne M, Beaudry P, Launay J-M, Laplanche J-L. A novel mutation (G114V) in the prion protein gene in a family with inherited prion disease. Neurology. 2005 Apr 26;64(8):1455–1457. PMID: 15851745

95. Liu Z, Jia L, Piao Y, Lu D, Wang F, Lv H, Lu Y, Jia J. Creutzfeldt-Jakob disease with PRNP G114V mutation in a Chinese family. Acta Neurol Scand. 2010 Jun;121(6):377–383. PMID: 20028338

96. Hsiao KK, Cass C, Schellenberg GD, Bird T, Devine-Gage E, Wisniewski H, Prusiner SB. A prion protein variant in a family with the telencephalic form of Gerstmann-Sträussler-Scheinker syndrome. Neurology. 1991 May;41(5):681–684. PMID: 1674116

97. Hinnell C, Coulthart MB, Jansen GH, Cashman NR, Lauzon J, Clark A, Costello F, White C, Midha R, Wiebe S, Furtado S. Gerstmann-Straussler-Scheinker disease due to a novel prion protein gene mutation. Neurology. 2011 Feb 1;76(5):485–487. PMID: 21282596

98. Panegyres PK, Toufexis K, Kakulas BA, Cernevakova L, Brown P, Ghetti B, Piccardo P, Dlouhy SR. A new PRNP mutation (G131V) associated with Gerstmann-Sträussler-Scheinker disease. Arch Neurol. 2001 Nov;58(11):1899–1902. PMID: 11709001

99. Jansen C, Parchi P, Capellari S, Strammiello R, Dopper EGP, van Swieten JC, Kamphorst W, Rozemuller AJM. A second case of Gerstmann-Sträussler-Scheinker disease linked to the G131V mutation in the prion protein gene in a Dutch patient. J Neuropathol Exp Neurol. 2011 Aug;70(8):698–702. PMID: 21760536

100. Hilton DA, Head MW, Singh VK, Bishop M, Ironside JW. Familial prion disease with a novel serine to isoleucine mutation at codon 132 of prion protein gene (PRNP). Neuropathol Appl Neurobiol. 2009 Feb;35(1):111–115. PMID: 19187063

101. Rowe DB, Lewis V, Needham M, Rodriguez M, Boyd A, McLean C, Roberts H, Masters CL, Collins SJ. Novel prion protein gene mutation presenting with subacute PSP-like syndrome. Neurology. 2007 Mar 13;68(11):868–870. PMID: 17353478

102. Kitamoto T, Iizuka R, Tateishi J. An amber mutation of prion protein in Gerstmann-Sträussler syndrome with mutant PrP plaques. Biochem Biophys Res Commun. 1993 Apr 30;192(2):525–531. PMID: 8097911

103. Kenny J, Woollacott I, Koriath C, Hosszu L, Adamson G, Rudge P, Rossor MN, Collinge J, Rohrer JD, Mead S. A novel prion protein variant in a patient with semantic dementia. J Neurol Neurosurg Psychiatry. 2017 Jun 1; PMID: 28572272

104. Fong JC, Rojas JC, Bang J, Legati A, Rankin KP, Forner S, Miller ZA, Karydas AM, Coppola G, Grouse CK, Ralph J, Miller BL, Geschwind MD. Genetic Prion Disease Caused by PRNP Q160X Mutation Presenting with an Orbitofrontal Syndrome, Cyclic Diarrhea, and Peripheral Neuropathy. J Alzheimers Dis JAD. 2017;55(1):249–258. PMCID: PMC5149415

105. Bommarito G, Cellerino M, Prada V, Venturi C, Capellari S, Cortelli P, Mancardi GL, Parchi P, Schenone A. A novel prion protein gene-truncating mutation causing autonomic neuropathy and diarrhea. Eur J Neurol. 2018 Aug;25(8):e91–e92. PMID: 29984897

106. Mead S, Gandhi S, Beck J, Caine D, Gajulapalli D, Gallujipali D, Carswell C, Hyare H, Joiner S, Ayling H, Lashley T, Linehan JM, Al-Doujaily H, Sharps B, Revesz T, Sandberg MK, Reilly MM, Koltzenburg M, Forbes A, Rudge P, Brandner S, Warren JD, Wadsworth JDF, Wood NW, Holton JL, Collinge J. A novel prion disease associated with diarrhea and autonomic neuropathy. N Engl J Med. 2013 Nov 14;369(20):1904–1914. PMCID: PMC3863770

107. Capellari S, Baiardi S, Rinaldi R, Bartoletti-Stella A, Graziano C, Piras S, Calandra-Buonaura G, D’Angelo R, Terziotti C, Lodi R, Donadio V, Pironi L, Cortelli P, Parchi P. Two novel PRNP truncating mutations broaden the spectrum of prion amyloidosis. Ann Clin Transl Neurol. 2018 Jun;5(6):777–783. PMCID: PMC5989776

108. Bishop MT, Pennington C, Heath CA, Will RG, Knight RSG. PRNP variation in UK sporadic and variant Creutzfeldt Jakob disease highlights genetic risk factors and a novel non-synonymous polymorphism. BMC Med Genet. 2009;10:146. PMCID: PMC2806268

109. Simpson M, Johanssen V, Boyd A, Klug G, Masters CL, Li Q-X, Pamphlett R, McLean C, Lewis V, Collins SJ. Unusual clinical and molecular-pathological profile of gerstmann-Sträussler-Scheinker disease associated with a novel PRNP mutation (V176G). JAMA Neurol. 2013 Sep 1;70(9):1180–1185. PMID: 23857164

110. Matsuzono K, Ikeda Y, Liu W, Kurata T, Deguchi S, Deguchi K, Abe K. A novel familial prion disease causing pan-autonomic-sensory neuropathy and cognitive impairment. Eur J Neurol Off J Eur Fed Neurol Soc. 2013 May;20(5):e67–69. PMID: 23577609

111. Medori R, Tritschler HJ, LeBlanc A, Villare F, Manetto V, Chen HY, Xue R, Leal S, Montagna P, Cortelli P. Fatal familial insomnia, a prion disease with a mutation at codon 178 of the prion protein gene. N Engl J Med. 1992 Feb 13;326(7):444–449. PMID: 1346338

112. Hitoshi S, Nagura H, Yamanouchi H, Kitamoto T. Double mutations at codon 180 and codon 232 of the PRNP gene in an apparently sporadic case of Creutzfeldt-Jakob disease. J Neurol Sci. 1993 Dec 15;120(2):208–212. PMID: 8138811

113. Nitrini R, Rosemberg S, Passos-Bueno MR, da Silva LS, Iughetti P, Papadopoulos M, Carrilho PM, Caramelli P, Albrecht S, Zatz M, LeBlanc A. Familial spongiform encephalopathy associated with a novel prion protein gene mutation. Ann Neurol. 1997 Aug;42(2):138–146. PMID: 9266722

114. Bütefisch CM, Gambetti P, Cervenakova L, Park KY, Hallett M, Goldfarb LG. Inherited prion encephalopathy associated with the novel PRNP H187R mutation: a clinical study. Neurology. 2000 Aug 22;55(4):517–522. PMID: 10953183

115. Collins S, Boyd A, Fletcher A, Byron K, Harper C, McLean CA, Masters CL. Novel prion protein gene mutation in an octogenarian with Creutzfeldt-Jakob disease. Arch Neurol. 2000 Jul;57(7):1058–1063. PMID: 10891990

116. Roeber S, Grasbon-Frodl E-M, Windl O, Krebs B, Xiang W, Vollmert C, Illig T, Schröter A, Arzberger T, Weber P, Zerr I, Kretzschmar HA. Evidence for a pathogenic role of different mutations at codon 188 of PRNP. PLOS One. 2008;3(5):e2147. PMCID: PMC2366066

117. Chen C, Shi Q, Zhou W, Zhang X-C, Dong J-H, Hu X-Q, Song X-N, Liu A-F, Tian C, Wang J-C, Gao C, Zhang J, Han J, Dong X-P. Clinical and familial characteristics of eight Chinese patients with T188K genetic Creutzfeldt-Jakob disease. Infect Genet Evol J Mol Epidemiol Evol Genet Infect Dis. 2013 Mar;14:120–124. PMID: 23261545

118. Shi Q, Zhou W, Chen C, Zhang B-Y, Xiao K, Zhang X-C, Shen X-J, Li Q, Deng L-Q, Dong J-H, Lin W-Q, Huang P, Jiang W-J, Lv J, Han J, Dong X-P. The Features of Genetic Prion Diseases Based on Chinese Surveillance Program. PLOS One. 2015;10(10):e0139552. PMCID: PMC4619501

119. Tartaglia MC, Thai JN, See T, Kuo A, Harbaugh R, Raudabaugh B, Cali I, Sattavat M, Sanchez H, DeArmond SJ, Geschwind MD. Pathologic evidence that the T188R mutation in PRNP is associated with prion disease. J Neuropathol Exp Neurol. 2010 Dec;69(12):1220–1227. PMCID: PMC3136530

120. Kotta K, Paspaltsis I, Bostantjopoulou S, Latsoudis H, Plaitakis A, Kazis D, Collinge J, Sklaviadis T. Novel mutation of the PRNP gene of a clinical CJD case. BMC Infect Dis. 2006;6:169. PMCID: PMC1693557

121. Zhang H, Wang M, Wu L, Zhang H, Jin T, Wu J, Sun L. Novel prion protein gene mutation at codon 196 (E196A) in a septuagenarian with Creutzfeldt-Jakob disease. J Clin Neurosci Off J Neurosurg Soc Australas. 2014 Jan;21(1):175–178. PMID: 23787189

122. Peoc’h K, Manivet P, Beaudry P, Attane F, Besson G, Hannequin D, Delasnerie-Lauprêtre N, Laplanche JL. Identification of three novel mutations (E196K, V203I, E211Q) in the prion protein gene (PRNP) in inherited prion diseases with Creutzfeldt-Jakob disease phenotype. Hum Mutat. 2000 May;15(5):482. PMID: 10790216

123. Dlouhy SR, Hsiao K, Farlow MR, Foroud T, Conneally PM, Johnson P, Prusiner SB, Hodes ME, Ghetti B. Linkage of the Indiana kindred of Gerstmann-Sträussler-Scheinker disease to the prion protein gene. Nat Genet. 1992 Apr;1(1):64–67. PMID: 1363809

124. Hsiao K, Dlouhy SR, Farlow MR, Cass C, Da Costa M, Conneally PM, Hodes ME, Ghetti B, Prusiner SB. Mutant prion proteins in Gerstmann-Sträussler-Scheinker disease with neurofibrillary tangles. Nat Genet. 1992 Apr;1(1):68–71. PMID: 1363810

125. Kim M-O, Cali I, Oehler A, Fong JC, Wong K, See T, Katz JS, Gambetti P, Bettcher BM, Dearmond SJ, Geschwind MD. Genetic CJD with a novel E200G mutation in the prion protein gene and comparison with E200K mutation cases. Acta Neuropathol Commun. 2013;1(1):80. PMCID: PMC3880091

126. Hsiao K, Meiner Z, Kahana E, Cass C, Kahana I, Avrahami D, Scarlato G, Abramsky O, Prusiner SB, Gabizon R. Mutation of the prion protein in Libyan Jews with Creutzfeldt-Jakob disease. N Engl J Med. 1991 Apr 18;324(16):1091–1097. PMID: 2008182

127. Mok TH, Koriath C, Jaunmuktane Z, Campbell T, Joiner S, Wadsworth JDF, Hosszu LLP, Brandner S, Parvez A, Truelsen TC, Lund EL, Saha R, Collinge J, Mead S. Evaluating the Causality of Novel Sequence Variants in the Prion Protein Gene by Example. Neurobiol Aging [Internet]. 2018 May 15 [cited 2018 May 22]; Available from: http://www.sciencedirect.com/science/article/pii/S0197458018301672

128. Heinemann U, Krasnianski A, Meissner B, Grasbon-Frodl EM, Kretzschmar HA, Zerr I. Novel PRNP mutation in a patient with a slow progressive dementia syndrome. Med Sci Monit Int Med J Exp Clin Res. 2008 May;14(5):CS41–43. PMID: 18443555

129. Piccardo P, Dlouhy SR, Lievens PM, Young K, Bird TD, Nochlin D, Dickson DW, Vinters HV, Zimmerman TR, Mackenzie IR, Kish SJ, Ang LC, De Carli C, Pocchiari M, Brown P, Gibbs CJ, Gajdusek DC, Bugiani O, Ironside J, Tagliavini F, Ghetti B. Phenotypic variability of Gerstmann-Sträussler-Scheinker disease is associated with prion protein heterogeneity. J Neuropathol Exp Neurol. 1998 Oct;57(10):979–988. PMID: 9786248

130. Komatsu J, Sakai K, Hamaguchi T, Sugiyama Y, Iwasa K, Yamada M. Creutzfeldt-Jakob disease associated with a V203I homozygous mutation in the prion protein gene. Prion. 2014 Sep 3;8(5):336–338. PMID: 25495585

131. Mastrianni JA, Iannicola C, Myers RM, DeArmond S, Prusiner SB. Mutation of the prion protein gene at codon 208 in familial Creutzfeldt-Jakob disease. Neurology. 1996 Nov;47(5):1305–1312. PMID: 8909447

132. Ripoll L, Laplanche JL, Salzmann M, Jouvet A, Planques B, Dussaucy M, Chatelain J, Beaudry P, Launay JM. A new point mutation in the prion protein gene at codon 210 in Creutzfeldt-Jakob disease. Neurology. 1993 Oct;43(10):1934–1938. PMID: 8105421

133. Pocchiari M, Salvatore M, Cutruzzolá F, Genuardi M, Allocatelli CT, Masullo C, Macchi G, Alemá G, Galgani S, Xi YG. A new point mutation of the prion protein gene in Creutzfeldt-Jakob disease. Ann Neurol. 1993 Dec;34(6):802–807. PMID: 7902693

134. Peoc’h K, Levavasseur E, Delmont E, De Simone A, Laffont-Proust I, Privat N, Chebaro Y, Chapuis C, Bedoucha P, Brandel J-P, Laquerriere A, Kemeny J-L, Hauw J-J, Borg M, Rezaei H, Derreumaux P, Laplanche J-L, Haïk S. Substitutions at residue 211 in the prion protein drive a switch between CJD and GSS syndrome, a new mechanism governing inherited neurodegenerative disorders. Hum Mol Genet. 2012 Dec 15;21(26):5417–5428. PMID: 22965875

135. Muñoz-Nieto M, Ramonet N, López-Gastón JI, Cuadrado-Corrales N, Calero O, Díaz-Hurtado M, Ipiens JR, Ramóny Cajal S, de Pedro-Cuesta J, Calero M. A novel mutation I215V in the PRNP gene associated with Creutzfeldt-Jakob and Alzheimer’s diseases in three patients with divergent clinical phenotypes. J Neurol. 2013 Jan;260(1):77–84. PMID: 22763467

136. Alzualde A, Indakoetxea B, Ferrer I, Moreno F, Barandiaran M, Gorostidi A, Estanga A, Ruiz I, Calero M, van Leeuwen FW, Atares B, Juste R, Rodriguez-Martínez AB, López de Munain A. A novel PRNP Y218N mutation in Gerstmann-Sträussler-Scheinker disease with neurofibrillary degeneration. J Neuropathol Exp Neurol. 2010 Aug;69(8):789–800. PMID: 20613639

137. Watts JC, Giles K, Serban A, Patel S, Oehler A, Bhardwaj S, Guan S, Greicius MD, Miller BL, DeArmond SJ, Geschwind MD, Prusiner SB. Modulation of Creutzfeldt-Jakob disease prion propagation by the A224V mutation. Ann Neurol. 2015 Oct;78(4):540–553. PMCID: PMC4711268

138. Jansen C, Parchi P, Capellari S, Vermeij AJ, Corrado P, Baas F, Strammiello R, van Gool WA, van Swieten JC, Rozemuller AJM. Prion protein amyloidosis with divergent phenotype associated with two novel nonsense mutations in PRNP. Acta Neuropathol (Berl). 2010 Feb;119(2):189–197. PMCID: PMC2808512

139. Bratosiewicz J, Barcikowska M, Cervenakowa L, Brown P, Gajdusek DC, Liberski PP. A new point mutation of the PRNP gene in Gerstmann-Sträussler-Scheinker case in Poland. Folia Neuropathol Assoc Pol Neuropathol Med Res Cent Pol Acad Sci. 2000;38(4):164–166. PMID: 11693719

140. Shepherd J, Cobbe SM, Ford I, Isles CG, Lorimer AR, MacFarlane PW, McKillop JH, Packard CJ. Prevention of coronary heart disease with pravastatin in men with hypercholesterolemia. West of Scotland Coronary Prevention Study Group. N Engl J Med. 1995 Nov 16;333(20):1301–1307. PMID: 7566020

141. Downs JR, Clearfield M, Weis S, Whitney E, Shapiro DR, Beere PA, Langendorfer A, Stein EA, Kruyer W, Gotto AM. Primary prevention of acute coronary events with lovastatin in men and women with average cholesterol levels: results of AFCAPS/TexCAPS. Air Force/Texas Coronary Atherosclerosis Prevention Study. JAMA. 1998 May 27;279(20):1615–1622. PMID: 9613910

142. Sabatine MS, Giugliano RP, Wiviott SD, Raal FJ, Blom DJ, Robinson J, Ballantyne CM, Somaratne R, Legg J, Wasserman SM, Scott R, Koren MJ, Stein EA, Open-Label Study of Long-Term Evaluation against LDL Cholesterol (OSLER) Investigators. Efficacy and safety of evolocumab in reducing lipids and cardiovascular events. N Engl J Med. 2015 Apr 16;372(16):1500–1509. PMID: 25773607

143. Ridker PM, Danielson E, Fonseca FAH, Genest J, Gotto AM, Kastelein JJP, Koenig W, Libby P, Lorenzatti AJ, MacFadyen JG, Nordestgaard BG, Shepherd J, Willerson JT, Glynn RJ, JUPITER Study Group. Rosuvastatin to prevent vascular events in men and women with elevated C-reactive protein. N Engl J Med. 2008 Nov 20;359(21):2195–2207. PMID: 18997196

144. ADAPT Research Group, Martin BK, Szekely C, Brandt J, Piantadosi S, Breitner JCS, Craft S, Evans D, Green R, Mullan M. Cognitive function over time in the Alzheimer’s Disease Anti-inflammatory Prevention Trial (ADAPT): results of a randomized, controlled trial of naproxen and celecoxib. Arch Neurol. 2008 Jul;65(7):896–905. PMCID: PMC2925195

145. Robinson JG, Farnier M, Krempf M, Bergeron J, Luc G, Averna M, Stroes ES, Langslet G, Raal FJ, El Shahawy M, Koren MJ, Lepor NE, Lorenzato C, Pordy R, Chaudhari U, Kastelein JJP, ODYSSEY LONG TERM Investigators. Efficacy and safety of alirocumab in reducing lipids and cardiovascular events. N Engl J Med. 2015 Apr 16;372(16):1489–1499. PMID: 25773378

146. Raal FJ, Santos RD, Blom DJ, Marais AD, Charng M-J, Cromwell WC, Lachmann RH, Gaudet D, Tan JL, Chasan-Taber S, Tribble DL, Flaim JD, Crooke ST. Mipomersen, an apolipoprotein B synthesis inhibitor, for lowering of LDL cholesterol concentrations in patients with homozygous familial hypercholesterolaemia: a randomised, double-blind, placebo-controlled trial. Lancet Lond Engl. 2010 Mar 20;375(9719):998–1006. PMID: 20227758

147. Rosas HD, Doros G, Gevorkian S, Malarick K, Reuter M, Coutu J-P, Triggs TD, Wilkens PJ, Matson W, Salat DH, Hersch SM. PRECREST: a phase II prevention and biomarker trial of creatine in at-risk Huntington disease. Neurology. 2014 Mar 11;82(10):850–857. PMCID: PMC3959748

148. Goldfarb LG, Petersen RB, Tabaton M, Brown P, LeBlanc AC, Montagna P, Cortelli P, Julien J, Vital C, Pendelbury WW. Fatal familial insomnia and familial Creutzfeldt-Jakob disease: disease phenotype determined by a DNA polymorphism. Science. 1992 Oct 30;258(5083):806–808. PMID: 1439789

149. Gabizon R, Rosenmann H, Meiner Z, Kahana I, Kahana E, Shugart Y, Ott J, Prusiner SB. Mutation and polymorphism of the prion protein gene in Libyan Jews with Creutzfeldt-Jakob disease (CJD). Am J Hum Genet. 1993 Oct;53(4):828–835. PMCID: PMC1682379

150. Webb TEF, Whittaker J, Collinge J, Mead S. Age of onset and death in inherited prion disease are heritable. Am J Med Genet Part B Neuropsychiatr Genet Off Publ Int Soc Psychiatr Genet. 2009 Jun 5;150B(4):496–501. PMID: 18729123

